# Cell cycle progression and stalked cell morphogenesis in *Caulobacter crescentus* are regulated independently by the flagellum

**DOI:** 10.64898/2026.11.02.763848

**Authors:** Chandler N. Hellenbrand, David M. Hershey

## Abstract

Bacteria colonize surfaces and form biofilms to survive in stressful environmental conditions. Many bacteria use an appendage called the flagellum to sense surface contact, and responding to these mechanical cues triggers rapid physiological changes to promote downstream attachment. Bacteria are known to secrete adhesins and alter their motility behavior to facilitate surface colonization, but these changes alone are insufficient to explain the breadth of adaptations observed in surface-attached cells. Disrupting flagellar assembly in the bacterium *Caulobacter crescentus* constitutively activates the surface sensing pathway, and this gain-of-function phenotype was previously used to identify a suite of *flagellar signaling suppressor (fss)* genes that promotes surface adhesion downstream of the flagellum. We show herein that a subset of *fss* genes also regulates the *C. crescentus* developmental program. Live-cell microscopy analysis of *fss* mutants identified independent sub-pathways that target stalked cell morphogenesis and cell cycle progression. We identify the predicted chemotaxis genes *fssD* and *fssE* as novel regulators of stalked cell morphogenesis and show that these genes act to stimulate the diguanylate cyclase PleD in flagellar assembly mutants. A separate pathway that includes the stator protein MotB and the recently characterized dNTPase FssC regulates entry into S phase downstream of the flagellum. We show that *motB* regulates cell cycle progression specifically in response to surface cues, and that stator-dependent cell cycle control occurs independently of c-di-GMP and CtrA. This work clarifies the downstream targets of the surface sensing network and establishes a comprehensive model for how surface sensing influences the *C. crescentus* developmental program.

**Importance:** Biofilm formation confers many advantages for bacterial cells, including stress resistance, protection from predators, and increased nutrient access. The majority of bacteria are predicted to reside in surface-attached communities, making biofilms central factors in human health, food and water management, and biogeochemical cycles. Understanding biofilm formation is crucial for manipulating these systems, but questions remain about how bacteria transition from a motile to a surface-attached state. This study identifies two genetic pathways that regulate growth and development after bacteria encounter surfaces and furthers our understanding of the cellular changes that occur during surface colonization.

## Introduction

Bacteria can colonize solid surfaces to form protected, multicellular communities called biofilms(1, 2). Surface colonization requires cells to recognize physical contact with target substrates and respond by activating signaling pathways that promote attachment. Motility appendages, such as flagella and pili, have a well-documented role in surface sensing(3–13). Flagella are complex trans-envelope machines that are assembled in a hierarchal manner(14, 15). Ion channels called stators bind to the inner cytoplasmic (C)-ring of the flagellum and use the proton motive force to rotate a flagellar filament, propelling the cell through its environment(16, 17). Current models for surface sensing implicate the stators as mechanosensors that recognize obstructions to filament rotation that occur during surface contact(5, 18). Likewise, surface contact is thought to disrupt the dynamic oscillation of type IV pili, and this tension is recognized as another physical surface sensing cue(19).

Mechanical signals sensed by flagella and pili activate diguanylate cyclase (DGC) enzymes that synthesize the second messenger c-di-GMP(7). c-di-GMP signaling directs cells to initiate biofilm formation and promotes maturation of the multicellular community. c-di-GMP binds effector proteins that alter gene expression, regulate motility and quorum sensing, promote virulence, and activate the secretion of polysaccharides(20–22). Together, these outputs of c-di-GMP regulation allow cells to develop into a collective biofilm in response to surface contact. This linear colonization sequence in which surface recognition activates c-di-GMP signaling to promote biofilm formation is present across a wide range of bacterial species, though there are exceptions to the critical role of c-di-GMP in surface attachment(23, 24).

The aquatic bacterium *Caulobacter crescentus* has emerged as an excellent model for studying surface adaptation(25). *C. crescentus* undergoes a complex developmental program with two distinct life cycle phases, the swarmer phase and the stalked phase(26, 27). Swarmer cells have type IV pili and a flagellum at one cell pole, and they differentiate into sessile stalked cells by shedding their motility appendages and extending the cell envelope to form a stalk (Fig. 1A). Stalked cells then divide asymmetrically to yield one of each cell type(28). The swarmer-to-stalked cell transition is regulated by a complex signaling network centered on c-di-GMP. The histidine kinases PleC and DivJ coordinate phosphorylation of the DGC PleD to activate c-di-GMP synthesis during the swarmer-stalked transition(29–31). PleD activates the morphogenic development of stalked cells by promoting ejection of the flagellum and stimulating stalk synthesis through the *shkA-tacA* pathway(30, 32–36). PleD also promotes cell cycle progression by deactivating the response regulator CtrA(37). CtrA acts both as a global transcription factor that regulates dozens of cell cycle-related genes(38) and a repressor of DNA replication initiation (39–41). c-di-GMP produced by PleD promotes deactivation and degradation of CtrA, allowing chromosome replication to proceed as swarmer cells differentiate into stalked cells.

**Figure 1:**
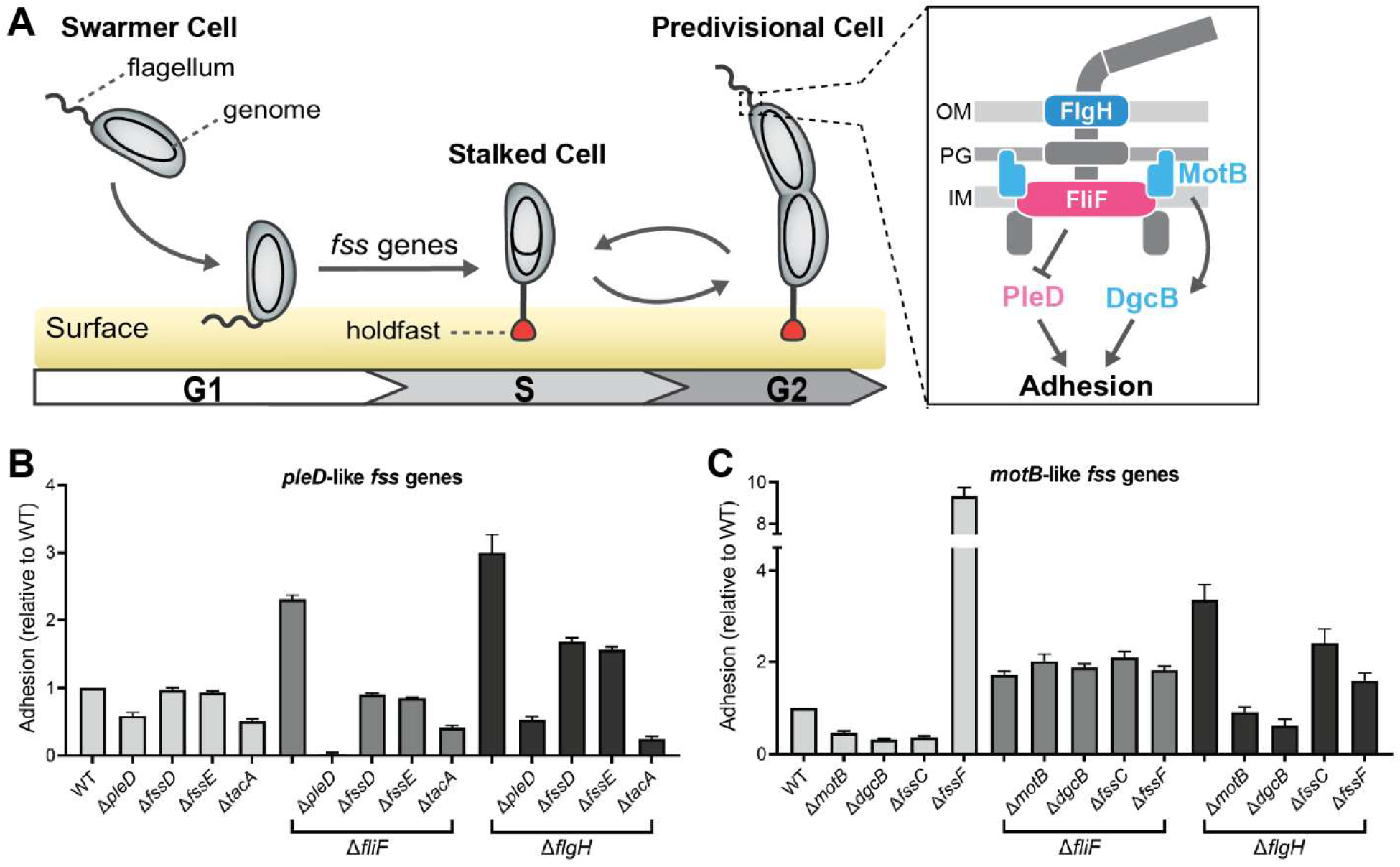
Adhesion in *C. crescentus* is regulated by *pleD* and *motB-*dependent pathways. A) Swarmer cells use their flagellum to sense surface contact and activate holdfast biosynthesis to mediate permanent surface attachment. Adhesion is regulated by two independent surface sensing pathways. The mechanical pathway (blue) contains the stators (MotB) and the diguanylate cyclase DgcB. The developmental pathway (pink) contains the diguanylate cyclase PleD. B) Deleting *fss* genes in the developmental pathway suppresses hyper-adhesion in both early and late flagellar assembly mutants (Δ*fliF* and Δ*flgH*, respectively). C). Deleting *fss* genes involved in the mechanical pathway suppresses hyper-adhesion only in the late flagellar assembly mutant Δ*flgH*. Error bars represent the standard deviation of the mean for three replicates.

Swarmer cells use their motility appendages to sense surface contact and stimulate the synthesis of an adhesin called the holdfast (Fig. 1A)(5, 19). Holdfast production can occur just minutes after surface contact(3), and it mediates irreversible attachment of *C. crescentus* cells to exogenous surfaces(42–44). Disrupting flagellar assembly stimulates holdfast production by activating the surface sensing pathway(18, 45). Deleting the late-stage flagellar assembly gene *flgH* results in partial synthesis of the flagellum, and the Δ*flgH* mutant is strongly hyper-adhesive due to an increase in holdfast production(18, 45). A suppressor screen identified a suite of surface sensing genes that contributes to hyper-adhesion in the Δ*flgH* mutant background(18). These *flagellar signaling suppressors* (*fss*) genes include the stator gene *motB* and the DGC-encoding genes *dgcB* and *pleD*. All three of these surface sensing genes regulate holdfast production, but they are differentially activated depending on the assembly state of the flagellum (Fig. 1A). Adhesion in late flagellar assembly mutants (*e.g*. Δ*flgH*) requires both a “mechanical” pathway dependent on *motB* and *dgcB* and a “developmental” pathway dependent on *pleD*, but adhesion in early flagellar assembly mutants (*e.g*. a Δ*fliF* mutant with no flagellar assembly) is entirely dependent on *pleD*(18). This bifurcated signaling topology underscores the inherent complexity of surface response pathways.

Related studies have shown that the type IV pilus mediates surface sensing in *C. crescentus*. Obstructing pilus retraction through chemical means stimulates holdfast production by mimicking disruptions to pilus dynamics thought to occur during surface contact(19). It was subsequently shown that activating type IV pilus-mediated surface sensing also stimulates DNA replication and stalked cell differentiation, leading to an abbreviated swarmer phase(9). The pilin subunit protein PilA seems to affect PleD phosphorylation by influencing PleC activity, but details of this signaling cascade remain vague(8). Nonetheless, these findings establish a precedent for surface cues regulating the *C. crescentus* developmental program. Whether flagellum-mediated surface sensing also influences stalked cell morphogenesis or cell cycle progression has remained unclear.

The breadth of surface sensing genes identified in the *fss* screen suggested a potential role of the flagellum in regulating *C. crescentus* growth and development. In addition to *pleD*, other developmental regulators, including *shkA* and *tacA*, contribute to hyper-adhesion in flagellar mutants(18). We also previously characterized a gene called *fssC* that regulates entry into the S phase of the cell cycle by degrading deoxynucleotide triphosphates (dNTPs)(46). Δ*fssC* cells have higher dNTP levels and show a CtrA-independent delay in the initiation of DNA replication(46). These findings prompted us to directly interrogate the impacts of flagellum-mediated surface sensing on the *C. crescentus* developmental program.

In this study, we analyze a panel of *fss* mutants (Table 1) and demonstrate that the flagellum regulates both stalked cell morphogenesis and cell cycle progression through independent pathways. We show that the predicted chemotaxis genes *fssD* and *fssE* regulate stalked cell morphogenesis through *pleD*, and that *fssD* and *fssE* specifically promote PleD activity when flagellar assembly is disrupted. A second pathway consisting of *motB* and *fssC* regulates entry into S phase independent of c-di-GMP and CtrA. We show that *motB* also controls cell cycle progression specifically when flagellar assembly is disrupted, and this effect can be recapitulated when load on the flagellum is increased by treating cells with methylcellulose. This work expands our understanding of surface responses that occur outside of canonical c-di-GMP signaling networks. Our results emphasize the complex effects of flagellum-mediated surface sensing on individual cells.

**Table 1:** *fss* genes analyzed in this study.

| ***fss* Gene Name** | **Description** | **CB15 Locus Tag** | **NA1000 Locus Tag** | **Reference(s)** |
|---|---|---|---|---|
| *motB* | Part of the stator complex. Required for adhesion through the mechanical pathway. | CC_1573 | CCNA_01644 | (5, 18) |
| *dgcB* | Diguanylate cyclase required for adhesion through the mechanical pathway. | CC_1850 | CCNA_01926 | (5, 18, 64) |
| *pleD* | Diguanylate cyclase required for adhesion through the developmental pathway. Regulates stalked cell morphogenesis. | CC_2462 | CCNA_02546 | (18, 34, 36) |
| *fssC* | dNTPase that regulates the G1-S phase transition. | CC_2008 | CCNA_02087 | (46) |
| *fssD* | CheYIV, shown herein to regulate stalked cell morphogenesis. | CC_3471 | CCNA_03585 | This work |
| *fssE* | CheRIII, shown herein to regulate stalked cell morphogenesis. | CC_3472 | CCNA_03586 | This work |
| *fssF* | FliNII, flagellar accessory protein required for chemotaxis. | CC_2175 | CCNA_02257 | (47) |
| *fssA* | Promotes stator function. See supplemental material for characterization. | CC_1064 | CCNA_01117 | (18) |
| *fssB* | Promotes stator function. See supplemental material for characterization. | CC_2058 | CCNA_02137 | (18) |

## Results

### Adhesion profiles of fss mutants phenocopy pleD or motB

Disrupting flagellar assembly stimulates adhesion through two distinct pathways: the mechanical (*motB*-dependent) pathway and the developmental (*pleD*-dependent) pathway (Fig. 1A)(18). Adhesion in late flagellar assembly mutants (Δ*flgH*) relies on both the mechanical and developmental pathways and can be suppressed by the deletion of either *motB* or *pleD* (Fig. 1BC). Adhesion in early flagellar assembly mutants (Δ*fliF*) is entirely dependent on *pleD* and is unaffected by genes in the mechanical pathway (Fig. 1BC)(18). We used crystal violet staining of adhesive cells to recapitulate the binning of *fss* genes (Table 1) into either the mechanical or developmental pathways based on their ability to promote hyper-adhesion in the Δ*fliF* mutant. The *fss* genes *dgcB*, *fssA*, *fssB*, *fssC*, and *fssF* follow a *motB-*dependent adhesion pattern. Their mutations specifically suppress hyper-adhesion in the Δ*flgH* background but have no effect in the Δ*fliF* background (Fig. 1C, S1)(18, 46, 47). The *fssA* and *fssB* genes encode putative flagellar components that have previously been shown to promote stator function(18). The Δ*fssA* and Δ*fssB* mutants resemble other stator mutants, and we will use Δ*motB* as a representative stator mutant throughout this work (Fig. S1). The *fssF* gene encodes a flagellar accessory protein required for chemotaxis(47). While deletion of *fssF* activates adhesion in the wild-type (WT) background, it follows a *motB-*dependent adhesion pattern in the flagellar mutant backgrounds (Fig. 1C).

In contrast, mutating *fss* genes known to control stalked cell development (*e.g*. *tacA*(33)) produces a *pleD-*like suppression pattern (Fig. 1B). We analyzed two uncharacterized genes (*fssD* and *fssE*) previously identified as flagellar surface sensing genes(18). The *fssD* and *fssE* genes form a two-gene operon, and each encode proteins with homology to chemotaxis signaling systems. FssD encodes a predicted CheY-like response regulator, and FssE encodes a predicted CheR-like methyltransferase (Fig. S2). Mutating *fssD* and *fssE* does not change adhesion levels in the WT background, but deleting either gene suppresses hyper-adhesion in both the Δ*fliF* and Δ*flgH* mutants (Fig. 1B). Expressing *fssD* and *fssE* at an ectopic location restored hyper-adhesion in the Δ*flgH* Δ*fssD* and Δ*flgH* Δ*fssE* double mutants (Fig. S3). The adhesion profiles of *fssD* and *fssE* clearly follow a *pleD-*dependent suppression pattern, suggesting that *fssD* and *fssE* regulate adhesion through the developmental pathway.

### Soft agar phenotypes reflect diverse roles for fss *genes*

We examined how various *fss* mutants spread in soft agar, as this assay provides an integrated readout of motility, chemotaxis, and developmental timing(18, 46–48). The Δ*flgH* and Δ*fssF* mutants do not spread through soft agar due to defects in motility and chemotaxis, respectively(47). A stator mutant (Δ*motB*) also does not spread through soft agar due to an inability to rotate the flagellar filament (Fig. 2AB)(18). The Δ*fssC* mutant shows a hyper-spreading phenotype, and we previously attributed this enhanced migration to an elongated swarmer phase of the cell cycle(46). The Δ*pleD* and Δ*dgcB* mutants have opposite soft agar phenotypes, despite both genes encoding DGCs. Deletion of *dgcB* causes 30% more spreading than WT, and the Δ*pleD* mutant spreads approximately 30% less than WT (Fig. 2B). The Δ*pleD* spreading phenotype is reportedly due to a defect in chemotaxis, though *pleD* mutants are notably hyper-motile in liquid(30, 34–36, 49). The Δ*fssD* and Δ*fssE* mutants retain the ability to spread through soft agar, confirming that both mutants are motile. However, they also spread less efficiently than WT in a manner that resembles the Δ*pleD* mutant (Fig. 2AB). Mutations that alter the duration of the swarmer phase often alter soft agar spreading behavior, and the frequency of such motility phenotypes among *fss* mutants suggested that flagellar surface sensing influences the swarmer-stalked transition.

**Figure 2:**
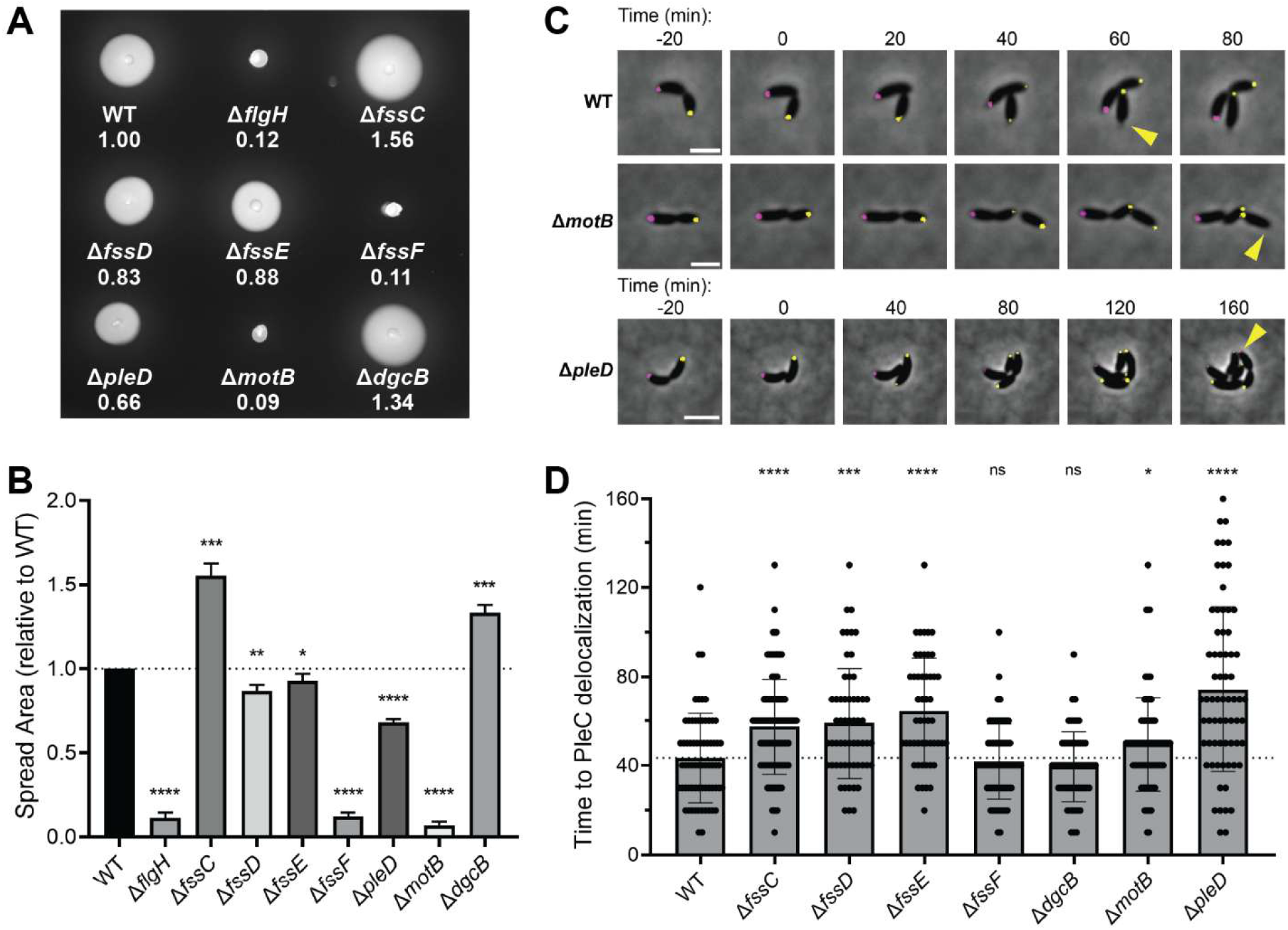
*fss* genes have diverse impacts on motility and development. A) Δ*fss* mutants display a diverse range of phenotypes in soft agar, indicating that the mutations impact motility, chemotaxis, and the cell cycle. B) Quantitative analysis of Δ*fss* mutant spreading in soft agar. Error bars represent the standard deviation of the mean for three replicates. C) Representative micrographs showing the localization of PleD-Venus and DivJ-mKate in WT, Δ*motB*, and Δ*pleD* cells over time. Yellow carets denote delocalization of PleC-Venus from the original swarmer pole. Scale bars represent 2 µm. D) Time required for PleC-Venus to delocalize in a panel of Δ*fss* mutants. The Δ*fssC*, Δ*fssD*, Δ*fssE*, Δ*motB*, and Δ*pleD* mutations delay PleC-Venus delocalization. Error bars represent the standard deviation of the mean for n > 50 cells. Unpaired t-tests were used to determine statistical significance. \**P* < 0.05, \*\**P* < 0.01, \*\*\**P* < 0.001, \*\*\*\**P* < 0.0001.

### The fss signaling network influences C. crescentus development

We directly investigated if mutating *fss* genes affects the *C. crescentus* developmental program. A reporter strain containing PleC and DivJ fused to different fluorescent proteins was previously developed to track the *C. crescentus* life cycle via live-cell microscopy (Fig. 2C)(46). We used this *pleC-venus divJ-mkate* strain to compare the time required for PleC-Venus to delocalize in newly divided swarmer cells among a panel of Δ*fss* mutants (Table 1)(Fig. 2D). This method was previously used to show that the Δ*fssC* mutant has an elongated swarmer phase(46). Indeed, deleting *fssC* delays PleC-Venus delocalization by almost 15 minutes compared to the WT strain (WT = 43 min, Δ*fssC =* 57 min). Most *fss* mutants also showed delays in PleC-Venus delocalization by at least an average of 7 minutes, including Δ*motB*, Δ*pleD*, Δ*fssD*, and Δ*fssE* mutants (Fig. 2D). The Δ*pleD* mutant had the most extreme phenotype with an average delay in PleC-Venus delocalization of 31 min, consistent with PleD’s critical role in stalked cell development. These findings indicate that *motB*, *fssD*, and *fssE* have newly described roles in regulating the *C. crescentus* developmental program. In contrast, the Δ*dgcB* and Δ*fssF* mutants did not show changes in the timing of PleC delocalization compared to WT (Δ*dgcB* = 39 min, Δ*fssF =* 41 min), suggesting that the roles of these genes may not extend to regulating development.

### fssD *and* fssE *promote stalked cell morphogenesis*

We next tested if any of the Δ*fss* mutants have defects in stalked cell morphogenesis. Differentiation into a stalked cell requires ejection of the flagellar filament and disassembly of the flagellar motor(14). We fluorescently tagged the flagellar C-ring protein FliG to visualize the assembly status of the flagellum (Fig. 3A)(47). Nearly all WT cells with fluorescent signal had a single FliG-Venus focus at either the pole of a swarmer cell or at the incipient swarmer pole of a predivisional cell. Only 1.4% of total cells in the WT population exhibited FliG-Venus foci at both cell poles (Fig. 3B). However, disrupting flagellar disassembly by deleting *pleD* increased the percentage of cells with two polar FliG-Venus foci to nearly 30% (Fig. 3B). This increase reflects a defect in flagellar disassembly, in which swarmer cells do not disassemble their flagellum before entering the predivisional phase of the life cycle(30, 34–36). The Δ*fssD* and Δ*fssE* mutants exhibited a similar increase in the percentage of cells with two polar FliG-Venus foci (27.1% and 28.7%, respectively), indicating that *fssD* and *fssE* are also required for efficient flagellar disassembly. None of the other *fss* mutants tested (Δ*fssC*, Δ*motB*, Δ*dgcB*, or Δ*fssF*) showed defects in flagellum disassembly (Fig. 3B), despite the Δ*fssC* and Δ*motB* mutants having an elongated swarmer phase (Fig. 2D).

**Figure 3:**
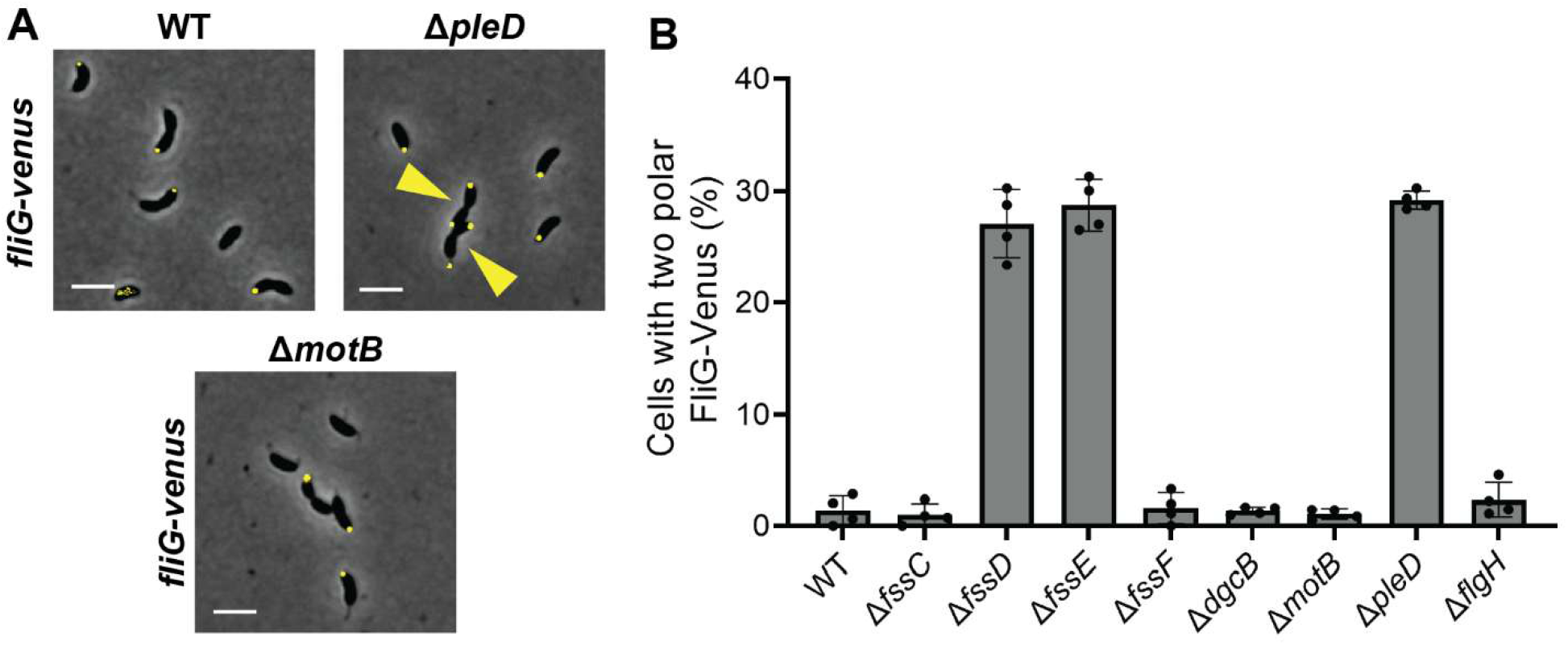
*fssD* and *fssE* promote flagellum disassembly. A) Micrographs showing FliG-Venus localization in WT, Δ*pleD*, and Δ*motB* cells. Yellow carets indicate cells with bipolar FliG-Venus localization. Scale bars represent 2 µm. B) Percentage of total cells with bipolar FliG-Venus localization in Δ*fss* mutant populations. The Δ*fssD*, Δ*fssE*, and Δ*pleD* strains have increased fractions of biflagellated cells. Error bars represent the standard deviation of the mean for four images containing n > 90 cells each.

Stalked cell morphogenesis is characterized by the extension of the cell envelope to form the stalk. Stalk synthesis is in part initiated by the bactofilin BacA, which localizes to the cell pole and recruits cell wall synthesis machinery (50, 51). We created a BacA-Venus fusion to track the timing of stalk synthesis with live-cell microscopy (Fig. 4A), and CV staining demonstrated that the *bacA-venus* allele was functional (Fig. S4). BacA localizes to the incipient stalked pole in an average of 22 min after cell division in WT cells. Deleting *pleD* delays BacA localization by 29 min, consistent with the disruption of stalked cell development (Fig. 4B). The Δ*fssD* and Δ*fssE* mutants again display a similar phenotype, with delays in BacA localization of 17 and 14 min, respectively. Deleting *fssC*, *motB*, *dgcB*, or *fssF* has no effect on the timing of BacA localization (Fig. 4B). Taken together, these data indicate that *pleD*, *fssD*, and *fssE* directly influence stalked cell morphogenesis, while *fssC*, *motB*, *dgcB*, and *fssF* do not.

**Figure 4:**
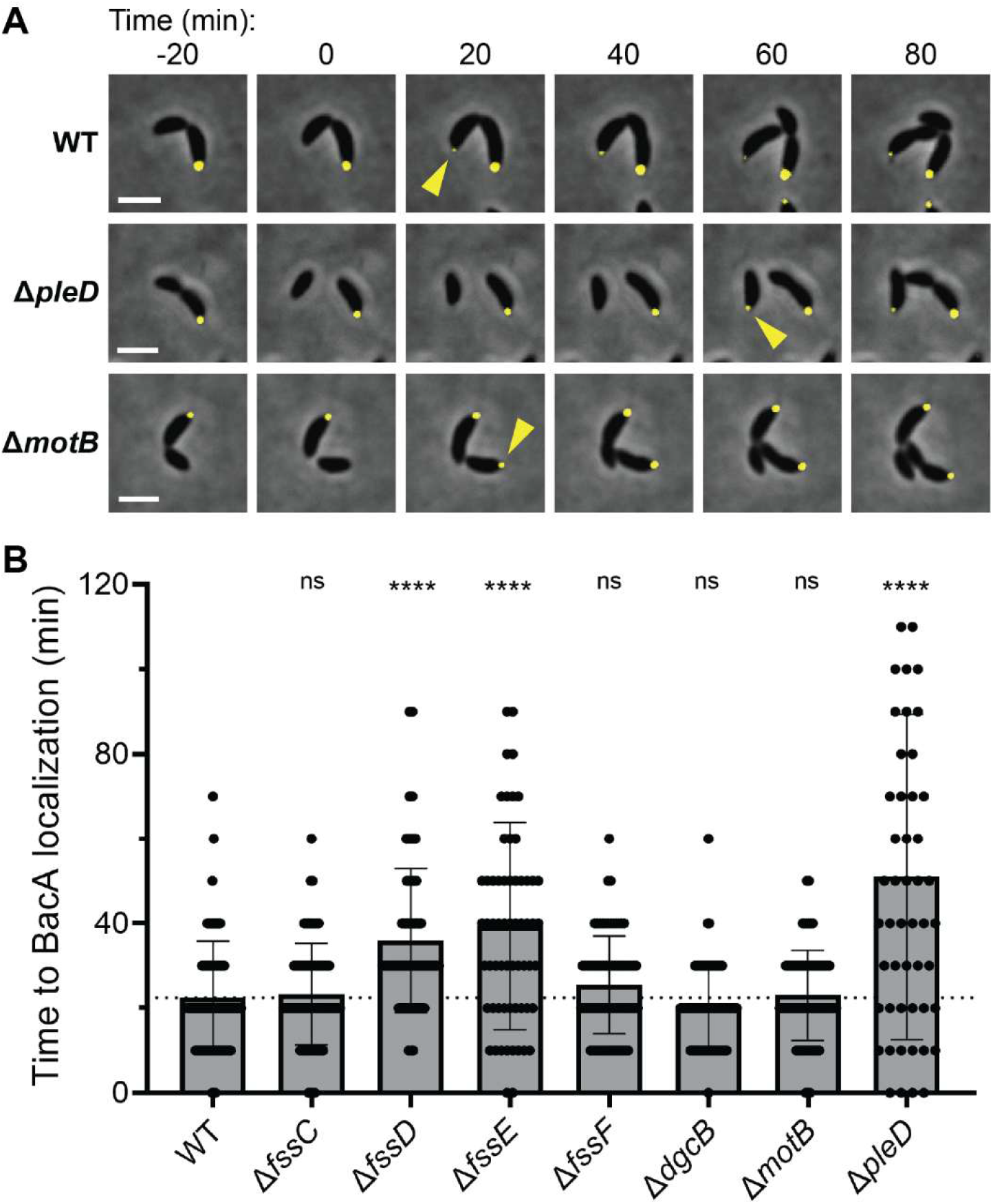
*fssD* and *fssE* promote stalk synthesis. A) Representative micrographs showing BacA-Venus localization in WT, Δ*pleD*, and Δ*motB* cells over time. Yellow carets denote BacA-Venus localization to new stalked poles. Scale bars represent 2 µm. B) Time required for BacA-Venus to localize in a panel of Δ*fss* mutants. Only the Δ*fssD*, Δ*fssE*, and Δ*pleD* mutants have delays in BacA-Venus localization compared to WT. Error bars represent the standard deviation of the mean for n > 50 cells. Unpaired t-tests were used to determine statistical significance. \*\*\*\**P* < 0.0001.

### pleD, fssD*, and* fssE *regulate stalked cell morphogenesis through the same pathway*

We combined the Δ*fssD,* Δ*fssE,* and Δ*pleD* mutations to determine if the three genes act in the same genetic pathway. *fssD* and *fssE* are adjacent genes in a predicted operon, and we constructed an Δ*fssDE* double mutant by deleting the entire coding sequence of the operon. The Δ*fssDE* mutation had indistinguishable adhesion from strains with single *fssD* or *fssE* deletions in the WT, Δ*fliF*, or Δ*flgH* backgrounds (Fig. 5A). *pleD* was also epistatic to *fssD* and *fssE*. The Δ*pleD* mutation suppressed adhesion more strongly than the Δ*fssDE* mutation in all genetic backgrounds tested, and combining the Δ*fssDE* Δ*pleD* deletions phenocopied the Δ*pleD* single deletions (Fig. 5A, 5D). These results suggest that *fssD* and *fssE* promote adhesion through the same pathway as *pleD*.

**Figure 5:**
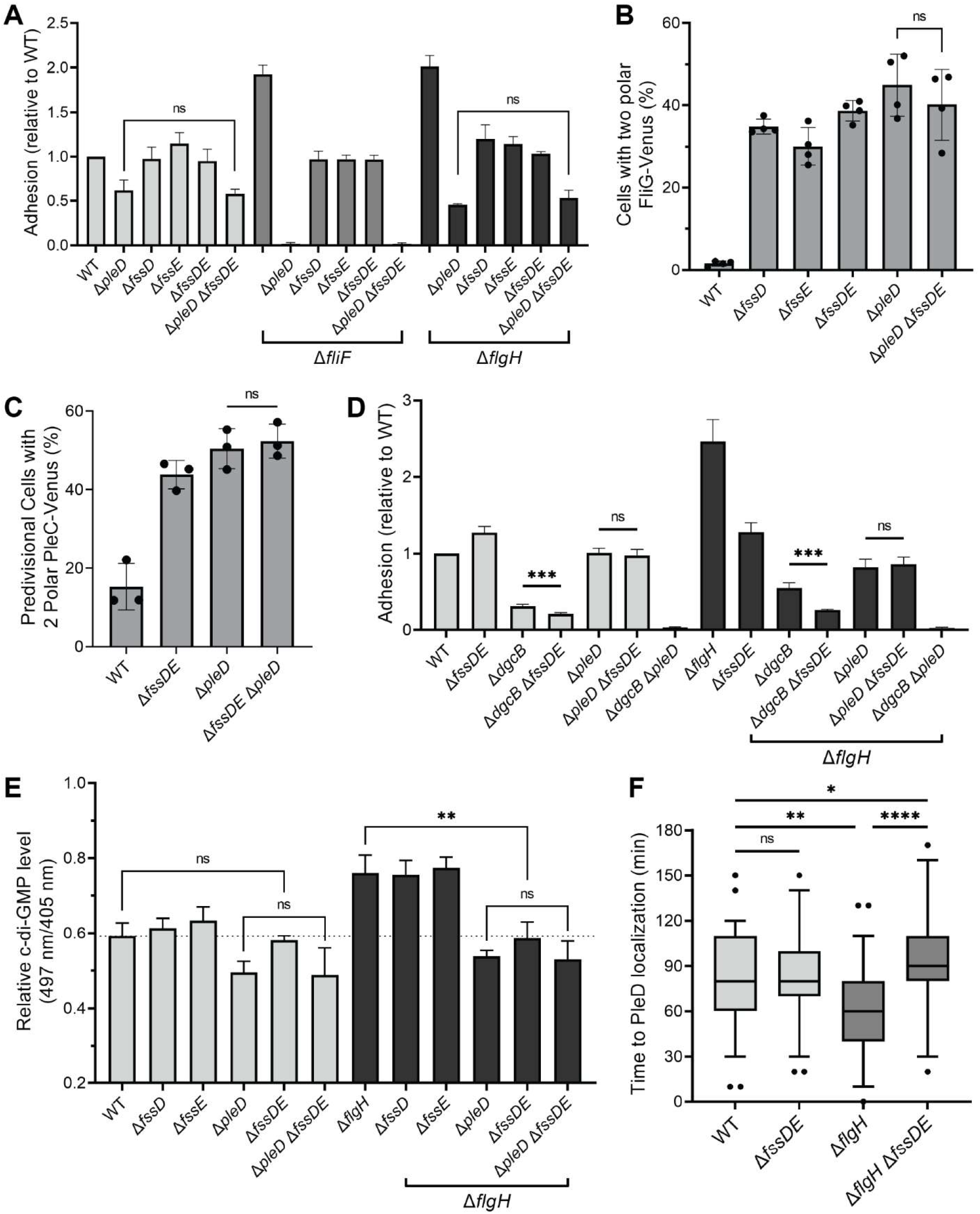
*fssD* and *fssE* regulate *pleD* activity in the Δ*flgH* mutant. A) Relative surface adhesion after deleting *pleD* and *fssDE* in WT, Δ*fliF*, Δ*flgH* genetic backgrounds. The Δ*pleD* and Δ*fssDE* phenotypes are not additive (*P* > 0.05). Error bars represent the standard deviation of the mean for three replicates. B) Percentage of cells with two polar FliG-Venus foci in Δ*pleD* and Δ*fssDE* populations. The Δ*pleD* and Δ*fssDE* mutations are not additive. Error bars represent the standard deviation of the mean for four images containing n > 93 cells each. C) Percentage of predivisional cells with two polar PleC-Venus foci in Δ*pleD* and Δ*fssDE* mutants. Increased bipolar PleC-Venus localization reflects a longer delay in PleC-Venus delocalization from the swarmer pole (Fig. S6). The Δ*pleD* and Δ*fssDE* mutations are not additive. Error bars represent the standard deviation of the mean for three replicates with n > 89 predivisional cells each. D) Adhesion phenotypes of Δ*fssDE* mutants are additive with Δ*dgcB*. Error bars represent the standard deviation of the mean for four replicates. E) Relative c-di-GMP levels measured with the ratiometric biosensor cdGreen(52). Fluorescence intensity was measured upon excitation at 497 and 405 nm and emission at 530 nm. Error bars represent the standard deviation of the mean for three replicates. F) Time required for PleD-Venus to localize to the stalked pole after deleting *fssDE* in WT and Δ*flgH* backgrounds. Error bars represent the 5-95 percentile with n > 50 cells. Unpaired t-tests were used to determine statistical significance in all panels. \**P* < 0.05, \*\**P* < 0.01, \*\*\**P* < 0.001, \*\*\*\**P* < 0.0001.

Adhesion in the WT and Δ*flgH* backgrounds is dependent on both *pleD* and *dgcB*. Combining the Δ*pleD* and Δ*dgcB* mutations causes additive effects that render cells non-adhesive (Fig. 5D). If *fssD* and *fssE* promote adhesion by working through *pleD*, then the adhesion defects of the Δ*dgcB* and Δ*fssDE* mutations should be additive. Indeed, the Δ*dgcB* Δ*fssDE* mutants are less adhesive than the Δ*dgcB* single mutants in both the WT and Δ*flgH* backgrounds (Fig. 5D), providing additional evidence that *fssD*, *fssE*, and *pleD* promote adhesion through the same pathway.

The developmental phenotypes associated with the Δ*pleD* and Δ*fssDE* mutations are also not additive. The percentage of cells with bipolar FliG-Venus localization in the Δ*pleD* mutant does not change when the Δ*fssDE* mutation is introduced (Fig. 5B). *pleD* is also completely epistatic to *fssD* and *fssE* in soft agar spreading assays (Fig. S5). We were unable to measure the timing of PleC-Venus delocalization via timelapse microscopy due to the extreme delay of the Δ*pleD* single mutant, so we instead analyzed the effects of *pleD* and *fssDE* on static PleC-Venus localization patterns (Fig. 5C). Some *C. crescentus* cells undergo a transient period in which PleC has not yet delocalized from the original swarmer pole but has already localized to the opposite, incipient swarmer pole. A longer delay in PleC delocalization extends the period in which cells have bipolar PleC localization (Fig. S6). As expected, approximately three times more predivisional cells show bipolar PleC localization in the Δ*pleD* and Δ*fssDE* mutants than in WT. The effects of *pleD* and *fssDE* were not additive in the Δ*pleD* Δ*fssDE* triple mutant (Fig. 5C), again indicating that *fssD* and *fssE* promote development through the same pathway as *pleD*.

### fssE *and* fssD *are likely upstream of* pleD

We measured c-di-GMP levels in Δ*fssD*, Δ*fssE*, and Δ*pleD* strains to investigate if *fssD* and *fssE* act upstream or downstream of *pleD*. We expressed the fluorescent protein biosensor cdGreen(52) from an ectopic location on the chromosome to measure c-di-GMP levels. Deleting *pleD* lowers c-di-GMP concentrations, but the Δ*fssD*, Δ*fssE,* and Δ*fssDE* mutations have no effect in the WT background (Fig. 5E). The Δ*pleD* Δ*fssDE* triple deletion mutant has similar c-di-GMP levels to the single Δ*pleD* mutant. In contrast, the Δ*flgH* mutant has elevated c-di-GMP levels that can be suppressed by deleting either *pleD* or *fssDE* (Fig. 5E). Deleting both Δ*pleD* and Δ*fssDE* in the Δ*flgH* background has indistinguishable c-di-GMP levels from the Δ*flgH* Δ*pleD* double mutant. This epistatic relationship places *fssD* and *fssE* upstream of *pleD*, but their effects on c-di-GMP levels are seen specifically when the surface sensing pathway is activated in the flagellar assembly mutant.

PleD localizes to the stalked cell pole when phosphorylated(31). We tracked the localization of a PleD-Venus fusion protein (Fig. S4) with live-cell microscopy. PleD localized to the stalked pole an average of 78 minutes after division in WT cells and 82 minutes in Δ*fssDE* cells, confirming that *fssD* and *fssE* have minimal effects on PleD activity in the WT background (Fig. 5F). However, activating the surface sensing pathway by deleting *flgH* caused earlier PleD localization, with foci appearing at the stalked cell pole in an average of 60 minutes. *fssD* and *fssE* were required for this accelerated PleD localization, as the combined Δ*flgH* Δ*fssDE* mutations delayed localization by 32 minutes compared to the single Δ*flgH* mutant (Fig. 5F). These results are consistent with the c-di-GMP measurements (Fig. 5E) and strongly indicate that *fssD* and *fssE* control *pleD* activity when surface sensing is activated.

### The stators promote early entry into S phase downstream of the flagellum

While the developmental delays observed in Δ*pleD*, Δ*fssD*, and Δ*fssE* mutants seem to derive from defects in stalked cell morphogenesis, deleting *fssC* or *motB* delayed PleC delocalization without disrupting flagellar disassembly or stalk synthesis (Fig. 2-4). We previously showed that *fssC* promotes the initiation of chromosome replication(46), and we reasoned that a subset of *fss* genes might affect development by targeting the G1-to-S phase transition. We used live-cell microscopy to measure the duration of the G1 phase by tracking the segregation of the chromosomal origin region with a fluorescent Venus-MipZ fusion protein (Fig. 6A)(46, 53). Consistent with previous results, the Δ*fssC* mutant showed a five minute delay in the duplication of MipZ compared to WT (Fig. 6B)(46). However, none of the other Δ*fss* mutants showed changes in the timing of MipZ duplication (Fig. 6B). Analyzing DNA content with flow cytometry (Fig. S7A) confirmed that the Δ*fssC* mutant has a clear increase in the proportion of cells with one copy of the chromosome, while the other Δ*fss* mutants do not (Fig. S7A).

**Figure 6:**
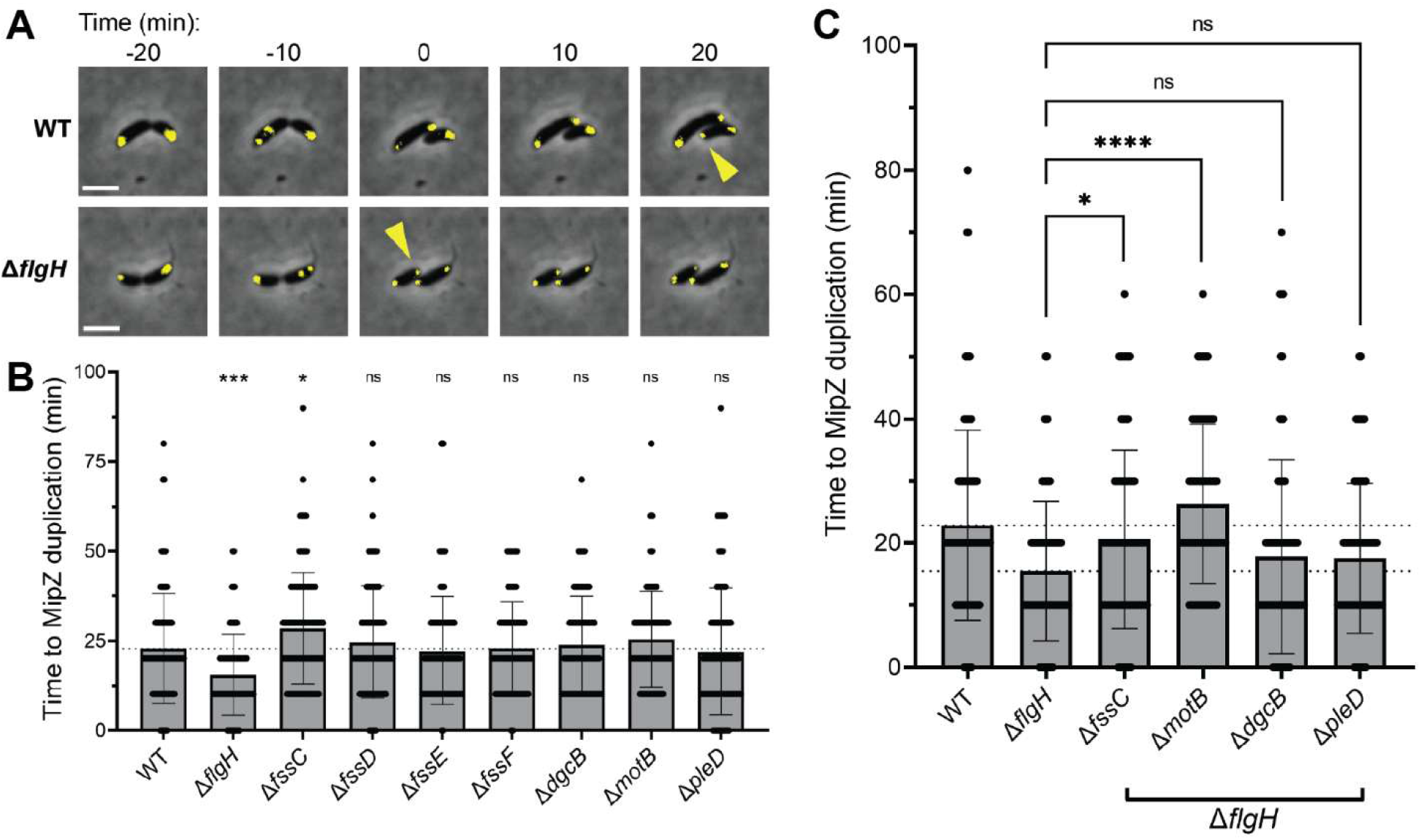
The stators promote entry into S phase downstream of the flagellum. A) Representative micrographs showing Venus-MipZ duplication in WT and Δ*flgH* cells over time. Yellow carets signify Venus-MipZ duplication. Scale bars represent 2 µm. B) Time required for Venus-MipZ to duplicate in Δ*flgH* and Δ*fss* mutants. Only the Δ*fssC* mutation delays Venus-MipZ duplication, consistent with FssC’s known role in promoting the G1-S phase transition. The Δ*flgH* mutant has earlier Venus-MipZ duplication. Error bars represent the standard deviation of the mean for n > 65 cells. C) Deletion of *fssC* and *motB* suppresses the Δ*flgH* mutant’s earlier entry into S phase. Error bars represent the standard deviation of the mean for n > 70 cells. Unpaired t-tests were used to determine statistical significance in all panels. \**P* < 0.05, \*\*\**P* < 0.001, \*\*\*\**P* < 0.0001.

Activating adhesion by deleting *flgH* shortened the G1 phase by eight minutes (Fig. 6B), and flow cytometry confirmed that the Δ*flgH* mutant has a lower proportion of cells in G1 phase (Fig. S7B). We sought to determine if any of the *fss* genes contribute to the accelerated G1-S phase transition in the Δ*flgH* background. The Δ*flgH* Δ*fssC* double mutant showed a 5-minute delay in MipZ duplication compared to the single Δ*flgH* mutant, which mirrors the effect of mutating *fssC* in the WT background (Fig. 6BC). However, deleting *motB* in the Δ*flgH* background delayed MipZ duplication by an average of 11 minutes (Fig. 6C). Because *motB* did not affect entry into S phase in WT cells (Fig. 6B), this result suggests that the stators become important regulators of the cell cycle when the surface sensing system is activated. The accelerated entry into S phase observed in the Δ*flgH* mutant appears to occur independently of c-di-GMP signaling, as deleting either *dgcB* or *pleD* in the Δ*flgH* background did not influence the timing of MipZ duplication (Fig. 6C). We used immunoblotting to investigate if the cell cycle regulator CtrA might be the mechanistic basis for the Δ*flgH* mutant’s early entry into S phase (Fig. S8-10). The Δ*flgH* strain indeed had earlier and more sustained depletion of CtrA, but this phenotype was not reversed by the deletion of *motB* (Fig. S8-10), indicating the stators promote entry into S phase independently of CtrA.

### *Cell cycle phenotypes of the Δ*fssC *and Δ*motB *mutants are not additive*

We investigated the relationship between *fssC* and *motB* in controlling *C. crescentus* development by constructing Δ*fssC* Δ*motB* double mutants(54). The Δ*fssC* and Δ*motB* mutations showed additive effects on surface adhesion in both the WT and Δ*flgH* backgrounds (Fig. 7A), suggesting that *fssC* and *motB* contribute to independent adhesion pathways. However, the cell cycle phenotypes of the Δ*fssC* and Δ*motB* mutants were not additive. PleC-Venus delocalization and Venus-MipZ duplication were not significantly different in the Δ*fssC* Δ*motB* double mutant compared to the single Δ*fssC* mutant (Fig. 7BC). In the Δ*flgH* background, the Δ*fssC* Δ*motB* double mutant’s delay in MipZ duplication is not significantly different from that in the Δ*motB* mutant (Fig. 7C). While the Δ*fssC* and Δ*motB* mutations do not have additive cell cycle phenotypes, there are discrepancies that suggest the two genes have a more complex relationship than simply being in the same versus independent pathways. *fssC* is epistatic to *motB* in the WT background (Fig. 7B), but *motB* is epistatic to *fssC* in the Δ*flgH* background (Fig. 7C). Together with the additive adhesion phenotype (Fig. 7A), we conclude that the *fssC* and *motB* pathways have some independent effects, but that their pathways converge on cell cycle regulation.

**Figure 7:**
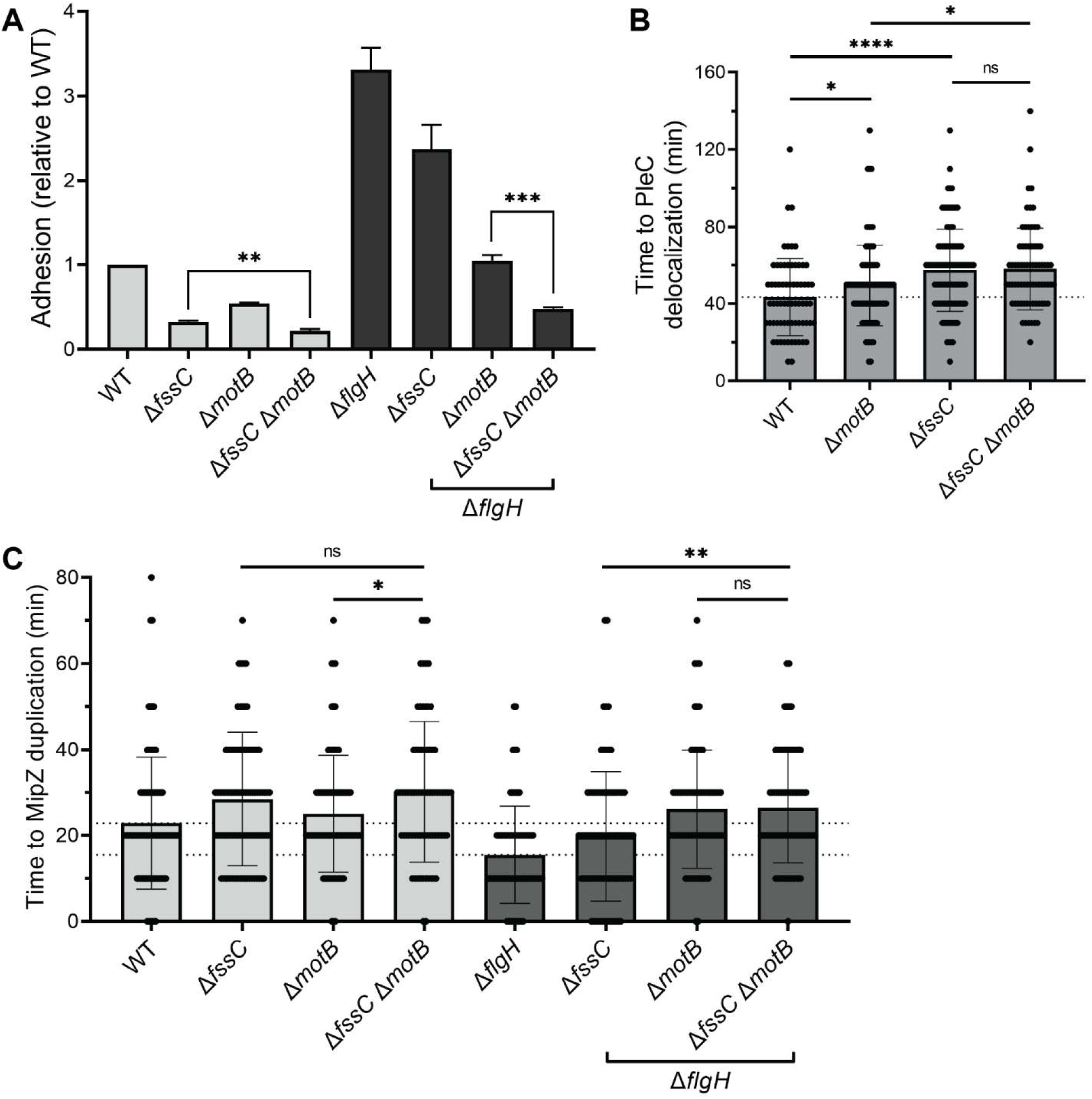
*fssC* and *motB* have related effects on the cell cycle. A) Relative adhesion after deleting *fssC* and *motB* in WT and Δ*flgH* genetic backgrounds. The Δ*fssC* and Δ*motB* phenotypes are additive. Error bars represent the standard deviation of the mean for three replicates. B) The Δ*fssC* and Δ*motB* mutants’ delays in PleC-Venus delocalization are not additive. Error bars represent the standard deviation of the mean for n > 68 cells each. C) The Δ*fssC* and Δ*motB* mutants’ delays in Venus-MipZ duplication are not additive in the WT or Δ*flgH* background. Error bars represent the standard deviation of the mean for n > 73 cells each. Unpaired t-tests were used to determine statistical significance in all panels. \**P* < 0.05, \*\**P* < 0.01, \*\*\**P* < 0.001, \*\*\*\**P* < 0.0001.

### motB *is required for surface-dependent cell cycle regulation*

We directly tested whether surface contact activates chromosome replication by adding methylcellulose to cells that were synchronized in the G1 phase and monitoring the duplication of Venus-MipZ. Methylcellulose activates surface sensing pathways by increasing the viscosity of liquid medium(54, 55). A higher fraction of WT cells entered S phase when methylcellulose was present (57% of cells by 60 minutes after synchronization) than in control cultures (37% of cells) without methylcellulose (Fig. 8A). We reasoned that if *fssC* or *motB* were involved in promoting cell cycle progression during surface sensing, then the fraction of cells entering S phase should not increase when these mutants are treated with methylcellulose. The Δ*fssC* mutant showed a similar surface response to WT cells, with 59% of cells entering S phase after 60 minutes in the methylcellulose condition compared to 49% without methylcellulose (Fig. 8A). In contrast, the Δ*motB* mutant completely lost the surface response. 35% and 34% of Δ*motB* cells entered S phase after 60 minutes with and without methylcellulose, respectively (Fig. 8A). These data directly demonstrate that *motB* promotes cell cycle progression during surface sensing but also suggest that *fssC* has additional activators independent of the flagellum.

**Figure 8:**
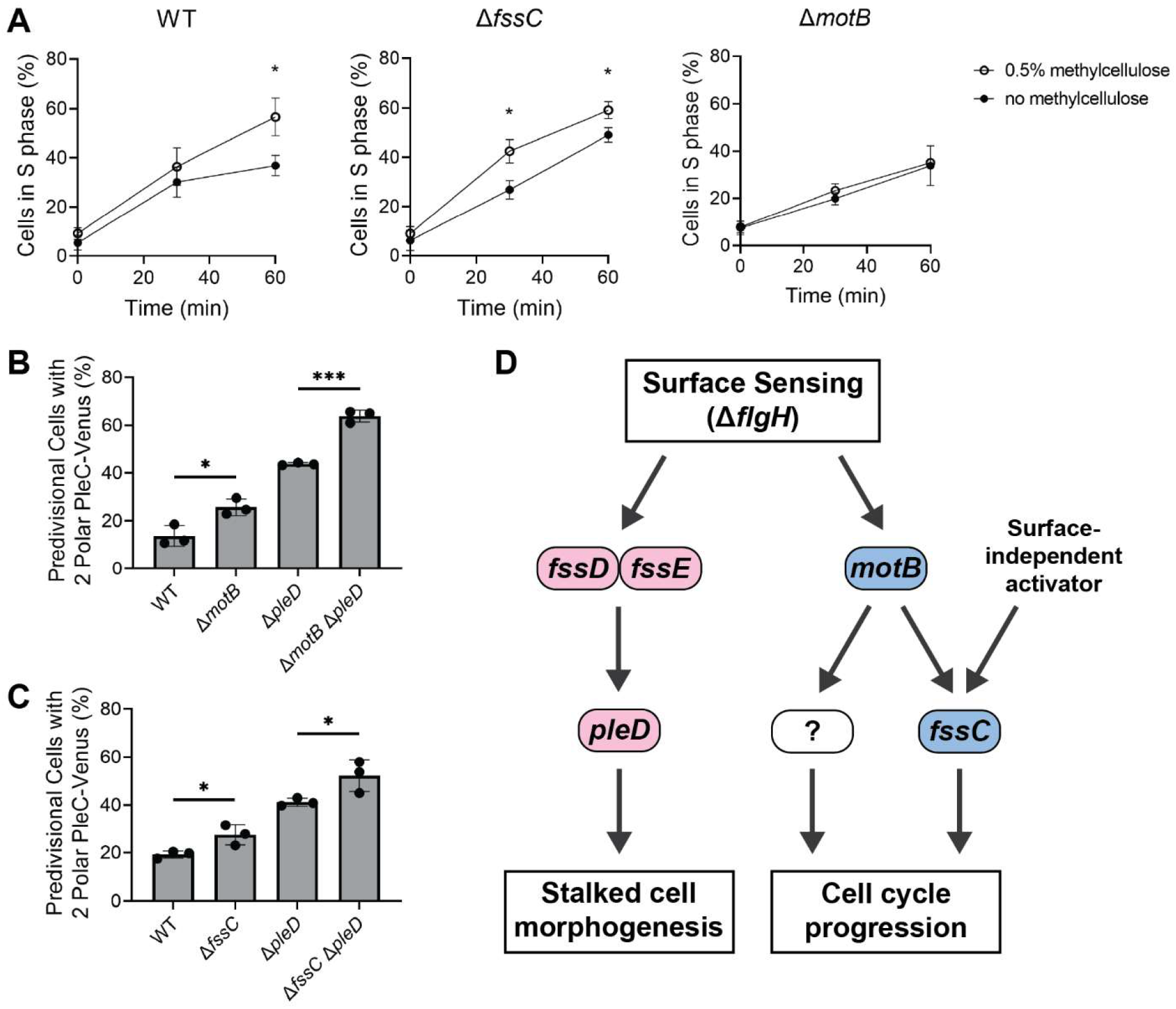
The flagellum regulates cell cycle progression and stalked cell morphogenesis independently. A) Synchronized swarmer cells were inoculated into liquid media with and without methylcellulose. The fraction of cells in S phase was determined by quantifying the number of cells with duplicated Venus-MipZ foci. Error bars represent the standard deviation of the mean for three replicates with n > 250 cells each. B) Percentage of predivisional cells with two polar PleC-Venus foci in Δ*pleD* and Δ*motB* mutants. Increased bipolar PleC-Venus localization is correlated with a longer delay in PleC-Venus delocalization from the swarmer pole (Fig. S6). The Δ*pleD* and Δ*motB* mutations are additive. Error bars represent the standard deviation of the mean for three replicates with n > 43 predivisional cells each. C) Percentage of predivisional cells with two polar PleC-Venus foci in Δ*pleD* and Δ*fssC* mutants. The Δ*pleD* and Δ*fssC* mutations are additive. Error bars represent the standard deviation of the mean for three replicates with n > 200 predivisional cells each. Unpaired t-tests were used to determine statistical significance in all panels. \**P* < 0.05, \*\*\**P* < 0.001. D) The flagellum promotes stalked cell morphogenesis (pink) and cell cycle progression (blue) through two independent pathways.

### The flagellum promotes cell cycle progression and stalk synthesis independently

Collectively, our results indicate that the *fss* network regulates *C. crescentus* development by targeting stalked cell morphogenesis through *fssD*, *fssE*, and *pleD* and cell cycle progression through *motB* and *fssC*. We constructed Δ*motB* Δ*pleD* and Δ*fssC* Δ*pleD* double mutants to determine if these pathways function independently. If the pathways are independent, then the mutants’ PleC localization phenotypes should be additive. We calculated the proportion of predivisional cells with bipolar PleC localization and found that both the Δ*motB* and Δ*fssC* mutations are additive with the Δ*pleD* mutation (Fig. 8BC). We conclude that stalked cell morphogenesis and cell cycle progression are regulated by the flagellum through separate, independent pathways (Fig. 8D).

## Discussion

Bacteria commonly form biofilms to construct favorable niches on surfaces. Understanding the molecular mechanisms of biofilm formation has implications for a number of health and environmental concerns(56–60). Surface colonization initiates when motile bacteria use extracellular appendages such as flagella and pili to sense surface contact (5, 19), triggering the downstream production of c-di-GMP by DGCs. c-di-GMP has long been known to induce physiological changes that promote the development of a surface-attached, multicellular community(20–22). In this study, we identify two surface sensing pathways that promote cellular differentiation and cell cycle progression outside of canonical c-di-GMP signaling. Our work highlights the complex effects of flagellum-mediated surface sensing and demonstrates how surface cues regulate the development and subcellular organization of individual cells.

We previously identified a suite of surface sensing genes that promotes the hyper-adhesion of flagellar assembly mutants (*flagellar signaling suppressor* (*fss*) genes) in *C. crescentus*. Two parallel pathways, one dependent on *motB*/*dgcB* and the other on *pleD*, independently activate holdfast biosynthesis when the flagellum is disrupted(Fig. 1A)(18). We expanded on our previous studies that have mapped the regulatory pathways for adhesion by examining the effects of *fss* genes (Table 1, Fig 1BC) on the *C. crescentus* developmental program. Tracking the delocalization of a PleC-Venus fusion protein in live cells identified widespread effects of the flagellar surface sensing system on *C. crescentus* development (Fig. 2C). All of the *fss* mutants tested, with the exception of Δ*fssF* and Δ*dgcB*, showed delays in PleC-Venus delocalization (Fig. 2D) that were caused by either a defect in stalked cell morphogenesis or in cell cycle progression. Further analysis of the flagellar assembly mutant Δ*flgH* showed heightened PleD activity and an accelerated G1-S phase transition (Fig. 5EF, 6), supporting the conclusion that the flagellum controls *C. crescentus* development through the *fss* signaling network.

We identified two previously uncharacterized genes (*fssD* and *fssE*) that show *pleD*-like adhesion patterns and are required for efficient PleC-Venus delocalization. The Δ*fssD* and Δ*fssE* mutants share numerous morphogenic defects with Δ*pleD* cells, including flagellar disassembly defects and stalk development delays (Fig. 3, 4). The Δ*pleD* and Δ*fssDE* mutations do not have additive effects in any phenotypic assay, indicating that all three genes regulate stalked cell development through the same pathway (Fig. 5, S5). In fact, *fssD* and *fssE* appear to act upstream of *pleD* in regulating adhesion and c-di-GMP production. Both genes are required for stimulating c-di-GMP production and efficient localization of PleD to the cell pole, but these phenotypic effects are seen only in the Δ*flgH* mutant and not in the WT background (Fig. 5EF). These results indicate that *fssD* and *fssE* control PleD activity specifically during surface sensing.

We propose that *fssD* and *fssE* are newly identified regulators of *pleD*. PleD activity is canonically regulated by PleC and DivJ, histidine kinases that coordinate PleD phosphorylation with the swarmer-stalked transition(29–31). While PleC is upstream of PleD, we show that the Δ*pleD* mutant has a delay in PleC delocalization (Fig. 2D). The exact sequence leading to PleC delocalization from the swarmer pole remains under debate (61, 62), but our results suggest that the mechanism is dependent on both stalked cell differentiation and entry into S phase. Previous work has also proposed the existence of an alternate PleD kinase that has yet to be identified(32). FssD and FssE are predicted chemotaxis proteins with CheY- and CheR-like domains, respectively (Fig. S2). Chemotaxis systems utilize histidine kinases to activate CheY-like proteins that bind the flagellar motor. It is possible that FssD and FssE are part of a broader chemotaxis-like system that includes a histidine kinase that phosphorylates PleD, but future work is necessary to investigate this hypothesis.

In contrast to *fssDE* and *pleD,* deleting *fssC* and *motB* delayed PleC delocalization without affecting flagellar disassembly or stalk synthesis (Fig. 2-4). The defects in Δ*fssC* and Δ*motB* cells appear instead to be caused by delayed cell cycle progression. *fssC* was previously found to promote the initiation of DNA replication(46), and here we showed that *motB* and *flgH* also influence the timing of the G1-S phase transition. MipZ duplication is accelerated in the Δ*flgH* mutant, and deleting *motB* delays MipZ duplication in the Δ*flgH* background but has no effect in WT cells(Fig. 6). This finding suggests that *motB* regulates cell cycle progression specifically in response to surface cues. Indeed, the Δ*motB* mutant lost the surface-dependent increase in cells entering S phase when grown in methylcellulose (Fig. 8A). The Δ*fssC* mutant, in contrast, retained the surface response seen in WT cells (Fig. 8A), indicating that *fssC* has additional activators independent of the flagellum (Fig. 8D).

The stators appear to regulate cell cycle progression independent of c-di-GMP and CtrA. *dgcB* and *pleD* were not required for the Δ*flgH* mutant’s early entry into S phase (Fig. 6C), and immunoblots indicated that depletion of CtrA is also not the mechanistic basis of stator-dependent cell cycle control (Fig. S8-10). Current surface sensing models implicate DgcB as a key player in transducing the mechanical signal sensed by the stators into a biochemical response that leads to adhesion(5). Our finding that the stators promote cell cycle progression independent of *dgcB* (Fig. 6C) suggests the presence of additional mechanisms for transducing the physical signal of surface contact. Epistasis analysis of the *fssC* and *motB* mutations showed that *motB* is likely upstream of *fssC* in the Δ*flgH* background, but both genes have some independent effects (Fig. 7). We propose that, in addition to *fssC*, an unknown regulator functions downstream of *motB* to promote the G1-S phase transition in response to surface cues (Fig. 8D).

While most of the *fss* mutants we analyzed in this study impacted *C. crescentus* growth and development, there were two exceptions. *fssF* encodes a recently characterized flagellar accessary protein required for chemotaxis(47), and *dgcB* encodes for a DGC that associates with the stators and promotes adhesion through the mechanical pathway (Fig. 1A)(5, 18). Neither *fssF* nor *dgcB*were found to influence PleC delocalization, flagellum disassembly, stalk synthesis, or cell cycle progression (Fig. 2-4, 6, S7), suggesting that their roles do not extend to regulating the *C. crescentus* developmental program. Previous work has identified *dgcB* as a developmental regulator (63, 64), but these studies focus on the effect of *dgcB* on motility and holdfast biosynthesis. Our results do not dispute these previous findings, as the Δ*dgcB* mutant indeed has decreased adhesion (Fig. 1C) and increased motility in soft agar (Fig. 2B).

This study adds to a growing consensus that the linear paradigm in which surface sensing activates c-di-GMP production to promote adhesion is insufficient to explain the wide range of surface-induced behaviors observed in bacteria. We identified two surface responses herein that add complexity to the accepted linear model of surface adaptation. First, the predicted chemotaxis genes *fssD* and *fssE* function upstream of PleD-dependent c-di-GMP signaling, indicating that c-di-GMP production is not always the initial response to surface contact. Second, *motB* appears to promote cell cycle progression independently of c-di-GMP. While c-di-GMP-independent surface responses are scarce in the literature(23, 24), our findings suggest it may be more prevalent.

Surface adaptation is often viewed through the lens of biofilm formation and the collective development of a multicellular community. Using *C. crescentus* as a model organism, it is apparent that surface contact also induces the subcellular reorganization of single cells. Such surface responses have also been reported in other bacterial species. A 1988 study in *Vibrio parahaemolyticus* found that flagellum-mediated surface sensing promotes swarmer cell differentiation in individual cells, and mutating the polar flagellum constitutively activates differentiation independent of surface contact(4). A more recent study in *Pseudomonas aeruginosa* found that surface contact triggers an asymmetric division program, in which one daughter cell is piliated and one is flagellated(65). Our work provides further evidence that surface contact influences growth and development at the single-cell level and reveals how the regulation of fundamental biological processes can change during surface colonization.

## Materials and Methods

### Bacterial strains, growth, and genetic manipulation

Strains used in this study are listed in Table S2. Plasmids were constructed with Gisbon assembly of PCR amplified inserts and plasmids linearized by restriction digest or PCR (listed in Table S1). Primer sequences are available upon request. *E. coli* was grown in LB medium at 37°C and supplemented with 50 µg/mL kanamycin when necessary. *C. crescentus* was grown in peptone yeast extract (PYE) medium or M2 minimal media supplemented with 0.15% xylose (M2X) at 30°C. All experiments were performed in PYE unless otherwise stated. PYE medium was supplemented with 5 µg/mL (for liquid medium) and 25 µg/mL (for solid medium) kanamycin when necessary. Liquid M2X medium was supplemented with 2 µg/mL kanamycin when necessary. Plasmids were transformed into *C. crescentus* by electroporation. Gene deletions and insertions were constructed with a two-step approach using *sacB-*based counterselection.

### Crystal violet-based adhesion assay

Strains were grown overnight in PYE and diluted to an OD_660_ of 0.5 before inoculating a 48-well plate. 2 µL of culture was used to inoculate 450 µL of M2X in each well. Well plates were incubated at 30°C shaking (200 rpm) for 18 hrs. Cultures were discarded and wells were rinsed with tap water. Adhered cells were stained by the addition of 0.5 mL of 0.01% crystal violet solution. Plates were incubated at room temperature for 15 minutes while shaking at 200 rpm before being washed again with tap water. 0.5 mL ethanol was then added to each well. Plates were incubated at room temperature for another 15 minutes while shaking before absorbances were read at 575 nm.

### Soft agar motility assay

Strains were grown overnight in PYE and diluted to an OD_660_ of 0.5 before plating. 2 µL were inoculated into PYE containing 0.25% agar. Soft agar plates were incubated at 30°C for at least 72 hrs before imaging. Spread areas were measured with ImageJ (66).

### Live-cell imaging of unsynchronized cells

Strains harboring *pleC-venus*/*divJ-mkate*, *venus-mipZ*, *bacA-venus*, or *pleD-venus* alleles were grown overnight in PYE. Strains were diluted in PYE to an OD_660_ of 0.05 and grown for 2 hrs before spotting 2.5 µL onto a 1.5% agarose pad made with PYE. Samples were incubated at 30°C for 1 hr before imaging. Microscopy was performed using a Nikon Ti-E inverted microscope equipped with an Orca Fusion BT digital CMOS camera (Hamamatsu). Fluorescence images were collected using a Lumencor Spectra III light engine light source and a YFP- and mCherry-specific filter set (Chroma). Images were collected every 10 min for 4 hrs. Image analysis was performed with MicrobeJ (67). All timelapse experiments were performed at least twice. A representative replicate with a minimum of 50 cells is shown.

### Quantifying FliG-Venus localization in cell populations

Strains were grown overnight in PYE and diluted to an OD_660_ of 0.05. Cultures were grown for 2 hrs before spotting 2 µL onto a 1% agarose pad made with water. Microscopy was performed with the same equipment described above. Four images were collected per strain containing at least 79 cells each.

### Quantifying PleC-Venus localization in cell populations

Strains were grown overnight in PYE and diluted to an OD_660_ of 0.05. Cultures were grown for 3-5 hrs before spotting 2 µL onto a 1% agarose pad made with water. Microscopy was performed with the same equipment described above. A minimum of 40 predivisional cells were counted over three images per replicate.

### Measuring relative c-di-GMP levels

Strains harboring pDH1619 at the *xyl* locus were grown in biological triplicate overnight in M2X supplemented with kanamycin. Cultures were diluted to an OD_660_ of 0.08 and grown for 6-8 hrs in M2X with kanamycin. 200 µL of each culture was transferred to 96-well plate in technical triplicate. Fluorescence intensity was measured upon excitation/emission at 497/530nm and normalized to the fluorescence intensity upon excitation/emission at 405/530nm (52).

### Flow cytometry

NA1000 strains were grown overnight in PYE and diluted to an OD_660_ of 0.05. Cultures were grown to an OD_660_ of 0.45 before fixing cells by the addition of cold 100% ethanol to a final concentration of 70%. Cells were stored at 4 °C for up to three days. Before analyzing, cells were pelleted by centrifugation at 4,000 x *g* for 3 min at 4 °C and resuspended in citrate buffer (55 mM, pH 9). Cells were diluted 1:10 in citrate buffer and left to incubate at room temperature for 3 hrs to let RNA degrade. Cells were stained with 0.25 µM SYTOX Green (Invitrogen). Cells were run on a CytoFLEX (Beckman Coulter) cytometer (Gains: FSC=143, SSC=74, FITC=250; Thresholds: SSC-A > 500 and FITC-A > 5000) with a 10 µL/min flow rate. Data was plotted in FlowJo version 10.10.00 (BD Life Sciences).

### Synchronization of NA1000 swarmer cells

NA1000 strains were grown overnight in PYE, diluted into M2X at an OD_660_ of 0.1, and grown for an additional 6-8 hrs. Cultures were diluted again into M2X such that the OD_660_ would be 0.5 at the desired time the following morning. Cells were harvested by centrifugation at 4,500 x *g* for 10 min at 4°C and resuspended in a 1:1 ratio of cold M2 salts and percoll. Swarmers were isolated by centrifugation at 7,000 x *g* for 20 min at 4°C. The bottom swarmer band was collected and washed once in M2 salts before use.

### Surface sensing experiment with methylcellulose

PYE containing 0.5% methylcellulose was prepared in advance by heating the media, adding methylcellulose, vortexing, and incubating at 4°C overnight. Synchronized swarmer cells were inoculated into PYE with and without methylcellulose at an OD_660_ of 0.04. Cultures were grown at room temperature and shaking at 55 RPM. 2 µL of cells were spotted onto 1% agarose pads made with water at desired timepoints. Microscopy was performed with the equipment described above. Two images were collected per replicate with a minimum of 80 cells each.

### Immunoblotting

Synchronized swarmer cells were inoculated into PYE at an OD_660_ of 0.1-0.2. Cells were normalized by OD_660_ and harvested by centrifugation at 16,000 x *g* for 3 min at 4°C. Cells were resuspended in tris-buffered saline with 1X Laemmli buffer and boiled for 10 min at 100°C. Lysates were frozen at −20°C before use. SDS-PAGE was performed with 12% acrylamide gels and proteins were transferred to a PVDF membrane in a semi-dry transfer apparatus (Trans-Blot Turbo, Biorad). Membranes were cut at the 20 kDa mark and the bottom portion was used as a loading control. Bottom portions were stained with 50% methanol, 10% acetic acid, and 0.1% Coomassie Brilliant Blue R-250 (Amresco) and destained with 50% methanol and 10% acetic acid. Top portions of the membranes were probed with a 1:10,000 dilution of anti-CtrA serum(68) followed by a 1:10,000 dilution of goat anti-rabbit (H+L)-horseradish peroxidase (Biorad). Blots were developed with Western Lightning chemiluminescence substrate (Revvity) and imaged with an iBright 1500 imager (Invitrogen).

## Data availability

Annotated plasmid maps, sequencing files, primer sequences, and cloning strategies for molecular manipulations are available upon request.

## Acknowledgements

We thank Dr. Briana Burton for providing access to a flow cytometer and Josie Reeve for assistance in making strains. This work was funded by the National Institutes of Health grant 1R35GM150652 to D.M.H. The funders had no role in study design, data collection and interpretation, or the decision to submit the work for publication.

## Supplemental Figures

**Fig. S1:**
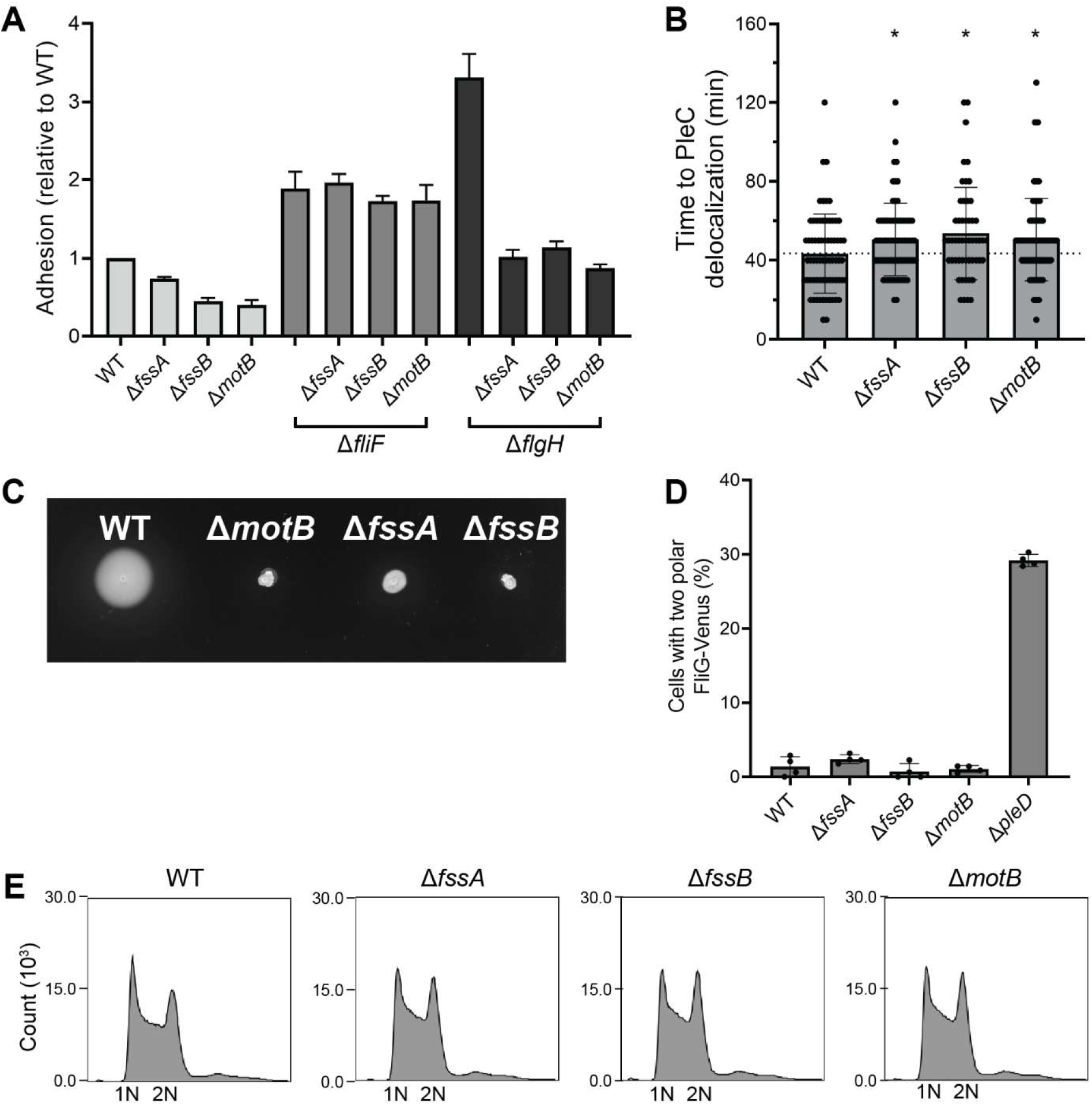
The Δ*fssA* and Δ*fssB* mutants are stator mutants and phenocopy Δ*motB*. A) The Δ*fssA* and Δ*fssB* mutants have a *motB*-dependent adhesion pattern. The mutations suppress adhesion in the WT and Δ*flgH* backgrounds but not in the Δ*fliF* background. Error bars represent the standard deviation of the mean for three replicates. B) The Δ*fssA*, Δ*fssB*, and Δ*motB* mutants have equivalent delays in PleC-Venus delocalization. Average time to PleC delocalization in WT = 43 min, Δ*fssA* = 51 min, Δ*fssB* = 54 min, Δ*motB* = 50 min. Error bars represent the standard deviation of the mean for n > 65 cells. Unpaired t-tests were used to determine statistical significance. \**P* < 0.05. C) The Δ*fssA* and Δ*fssB* mutants have spreading defects in soft agar similar to the Δ*motB* mutant. D) Δ*fssA*, Δ*fssB*, and Δ*motB* mutations do not impact flagellum disassembly. The Δ*pleD* mutant is shown as a positive control. Error bars represent the standard deviation of the mean for four images containing n > 79 cells each. E) DNA content analyzed by flow cytometry. The Δ*fssA*, Δ*fssB*, and Δ*motB* mutants do not display a delay in the G1-S phase transition.

**Fig. S2:**
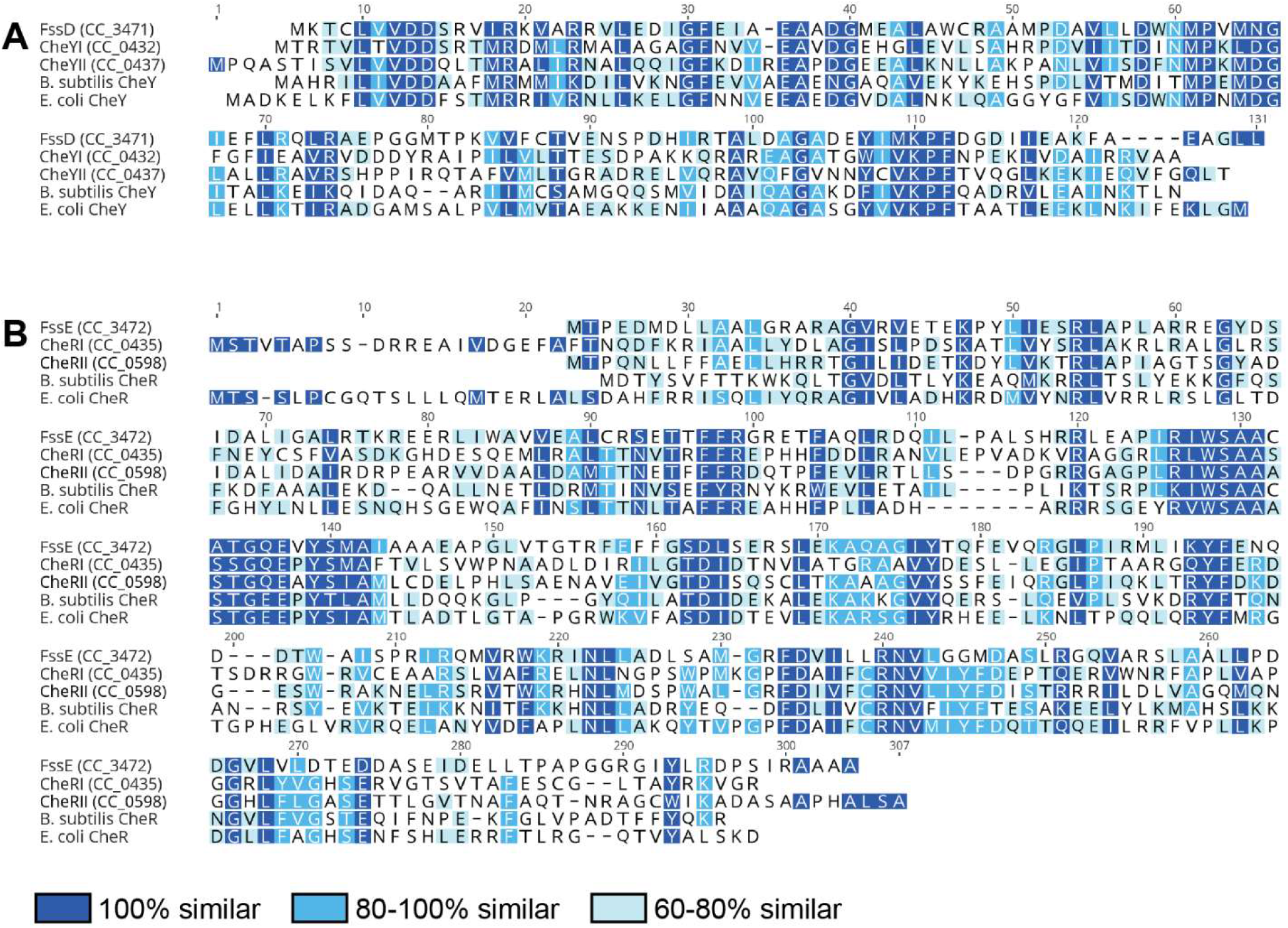
FssD and FssE are predicted CheY and CheR-like proteins. A) Clustal Omega alignment of the FssD protein sequence to other CheY proteins in *C. crescentus*, *Bacillus subtilis*, and *Escherichia coli*. B) Clustal Omega alignment of the FssE protein sequence to other CheR proteins in *C. crescentus*, *Bacillus subtilis*, and *Escherichia coli*.

**Fig. S3:**
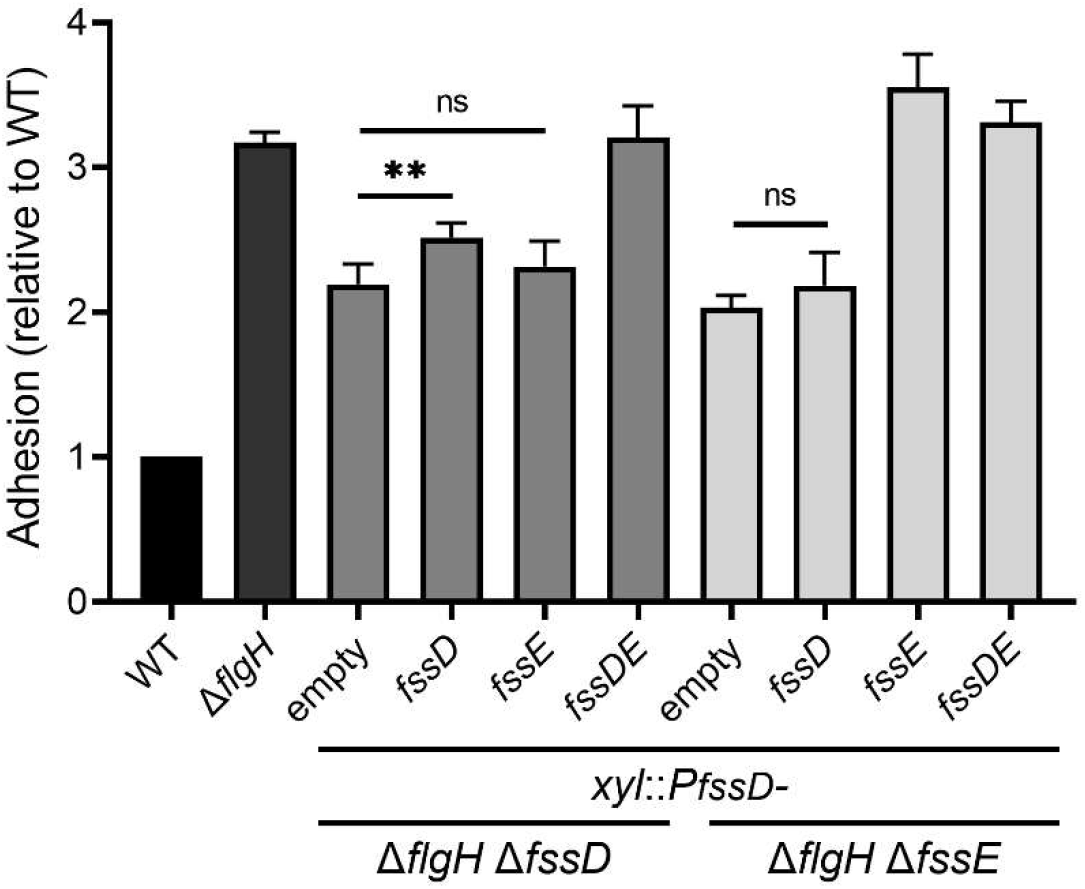
Expressing *fssD* and *fssE* at an ectopic locus restores adhesion in Δ*flgH* Δ*fssD* and Δ*flgH* Δ*fssE* mutant backgrounds. *fssD* and *fssE* were integrated at the *xyl* locus and expressed using their native promoter from the *fssDE* operon (*P_fssD_*, first 250 bp upstream of *fssD* start codon). The *P_fssD_*-*fssE* construct included the last 80 nucleotides of the *fssD* coding region to account for a potential ribosome binding site. Adhesion in the Δ*flgH* Δ*fssD* mutant was not restored by expressing only the promoter or *fssE*. Expressing *fssD* partially restored adhesion, and expressing the entire *fssDE* operon fully complemented the mutant. Adhesion in the Δ*flgH* Δ*fssE* mutant was not restored by expressing only the promoter or *fssD*. Expressing *fssE* and the *fssDE* operon fully restored adhesion. This genetic complementation rules out the possibility of polar effects in the Δ*fssD* and Δ*fssE* mutants. Error bars represent the standard deviation of the mean for four replicates. Statistical comparisons were made with unpaired t-tests. \*\**P* < 0.01

**Fig. S4:**
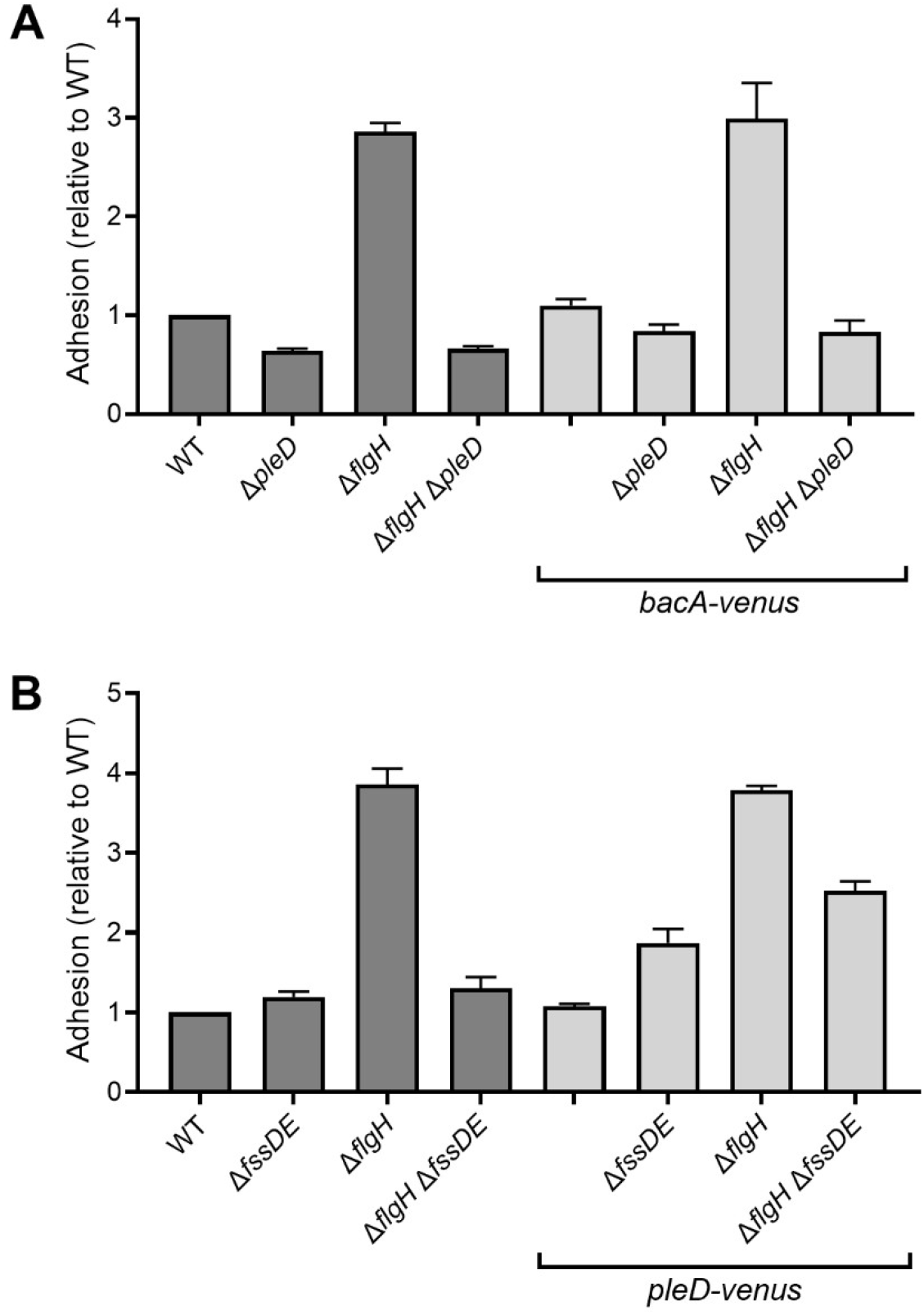
The *bacA-venus* and *pleD-venus* alleles are functional. A) The replacement of *bacA* with *bacA-venus* at its native locus does not affect adhesion levels in Δ*pleD* or Δ*flgH* strains. B) The replacement of *pleD* with *pleD-venus* at its native locus has minimal effects on adhesion in Δ*fssDE* and Δ*flgH* strains. There is a slight increase in adhesion observed in strains with *fssDE* deleted, but overall trends are consistent with the untagged strains. Error bars represent the standard deviation of the mean for three replicates.

**Fig. S5:**
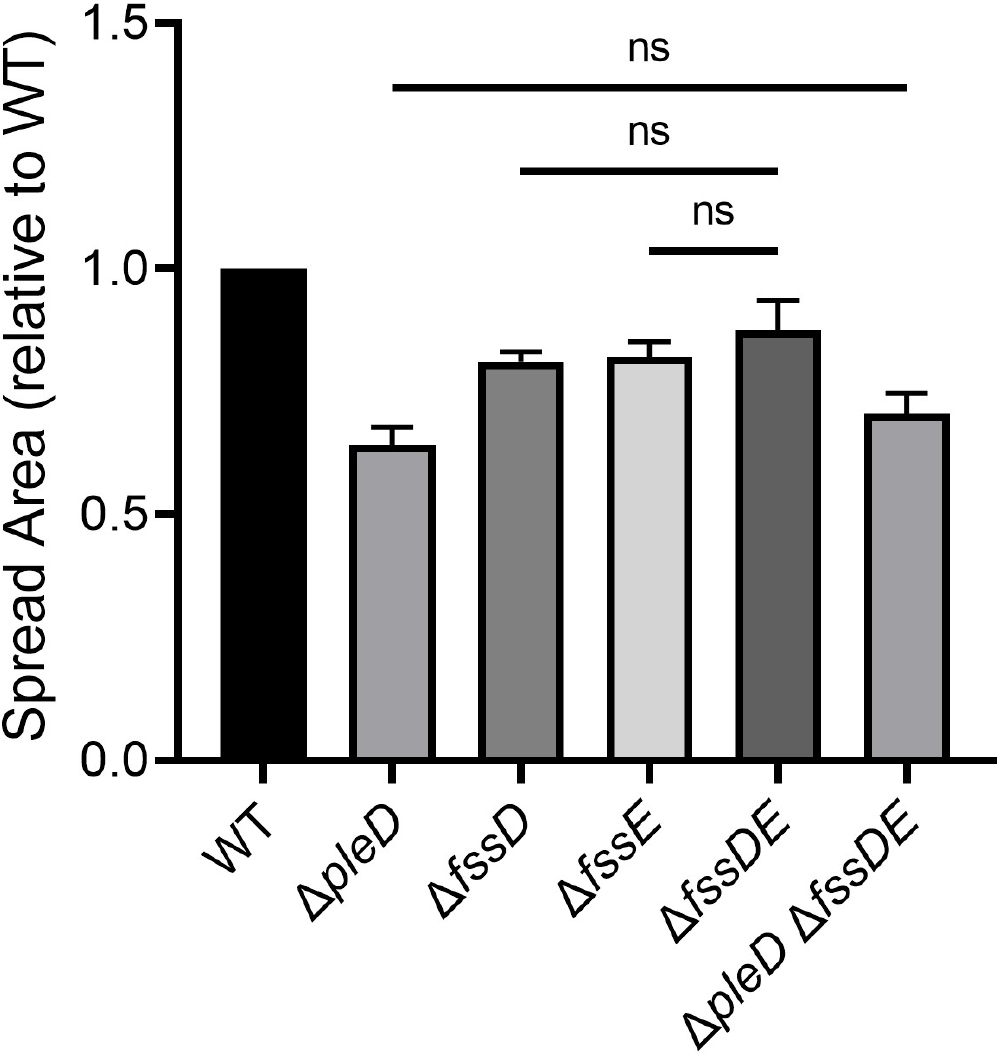
The Δ*pleD* and Δ*fssDE* soft agar phenotypes are not additive. *pleD* is epistatic to *fssDE* on soft agar. Error bars represent the standard deviation of the mean for four replicates. Statistical comparisons were made with unpaired t-tests.

**Fig. S6:**
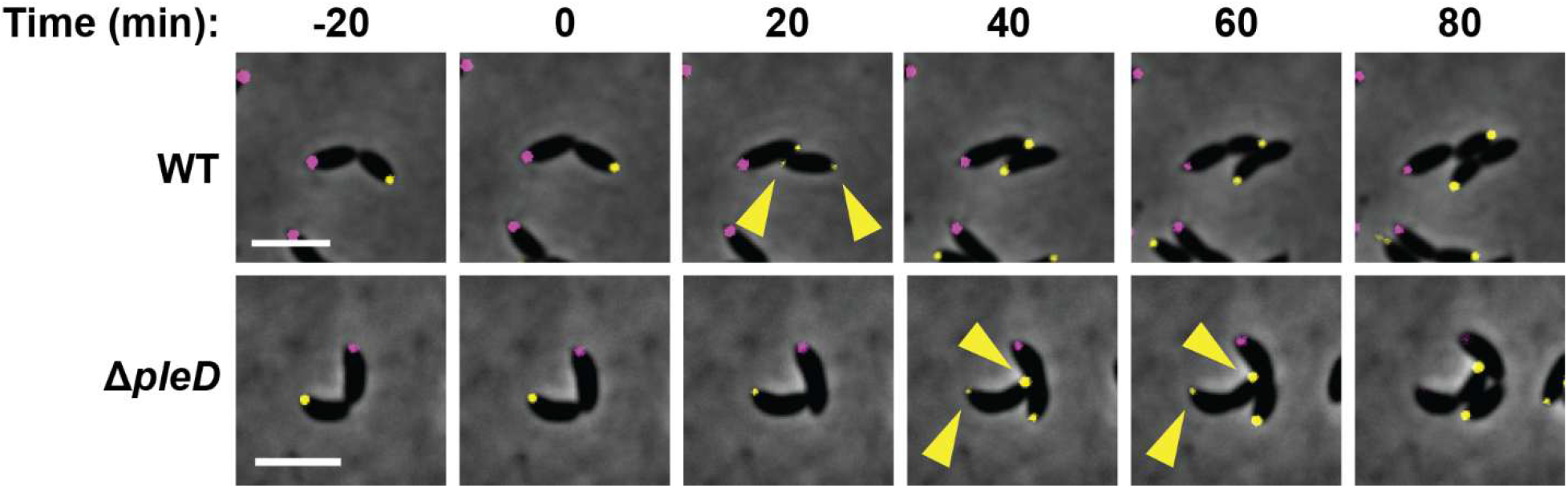
Delayed PleC delocalization from the original swarmer pole leads to a longer period of bipolar PleC localization. Representative micrographs showing transient bipolar PleC-Venus localization in WT and Δ*pleD* cells (denoted by yellow carets). DivJ-mKate marks the stalked poles. Scale bars represent 2 µm.

**Fig. S7:**
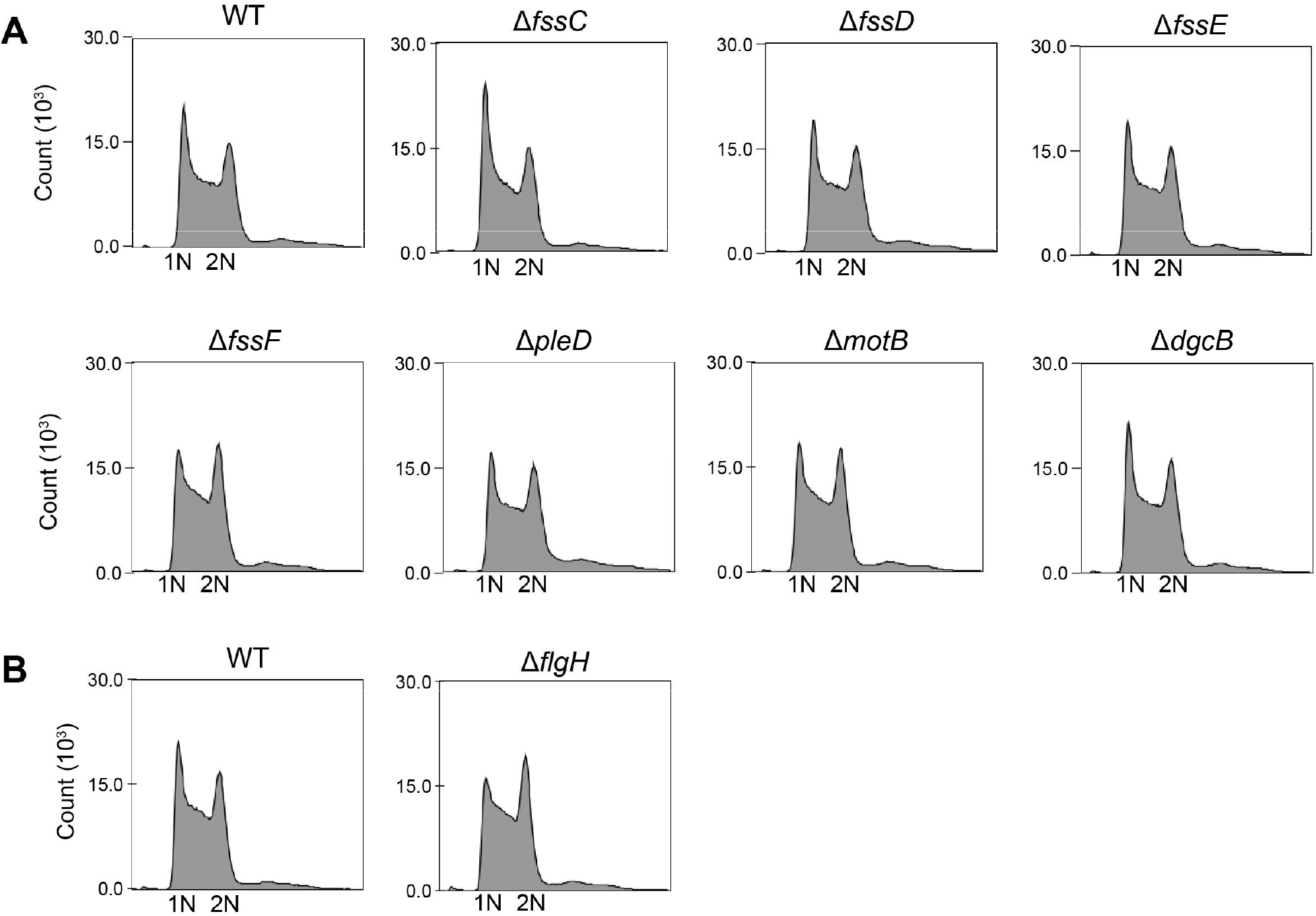
Analyzing DNA content by flow cytometry supports the Venus-MipZ duplication phenotypes. A) Of the Δ*fss* mutants tested, only the Δ*fssC* mutant has an increase in cells with one copy of the chromosome (1N). This is consistent with the known role of *fssC* in regulating the G1-S phase transition and the data presented in Fig. 6B, in which only the Δ*fssC* mutant has a delay in Venus-MipZ duplication compared to WT. The other Δ*fss* mutants do not have delays in the G1-S phase transition are within the normal range seen for WT populations. B) The Δ*flgH* mutant has an increase in cells with two copies of the chromosome (2N). This is again consistent with the data presented in Fig. 6B, in which the Δ*flgH* mutant has earlier Venus-MipZ duplication compared to WT. Samples in each panel were run on the same day.

**Fig. S8:**
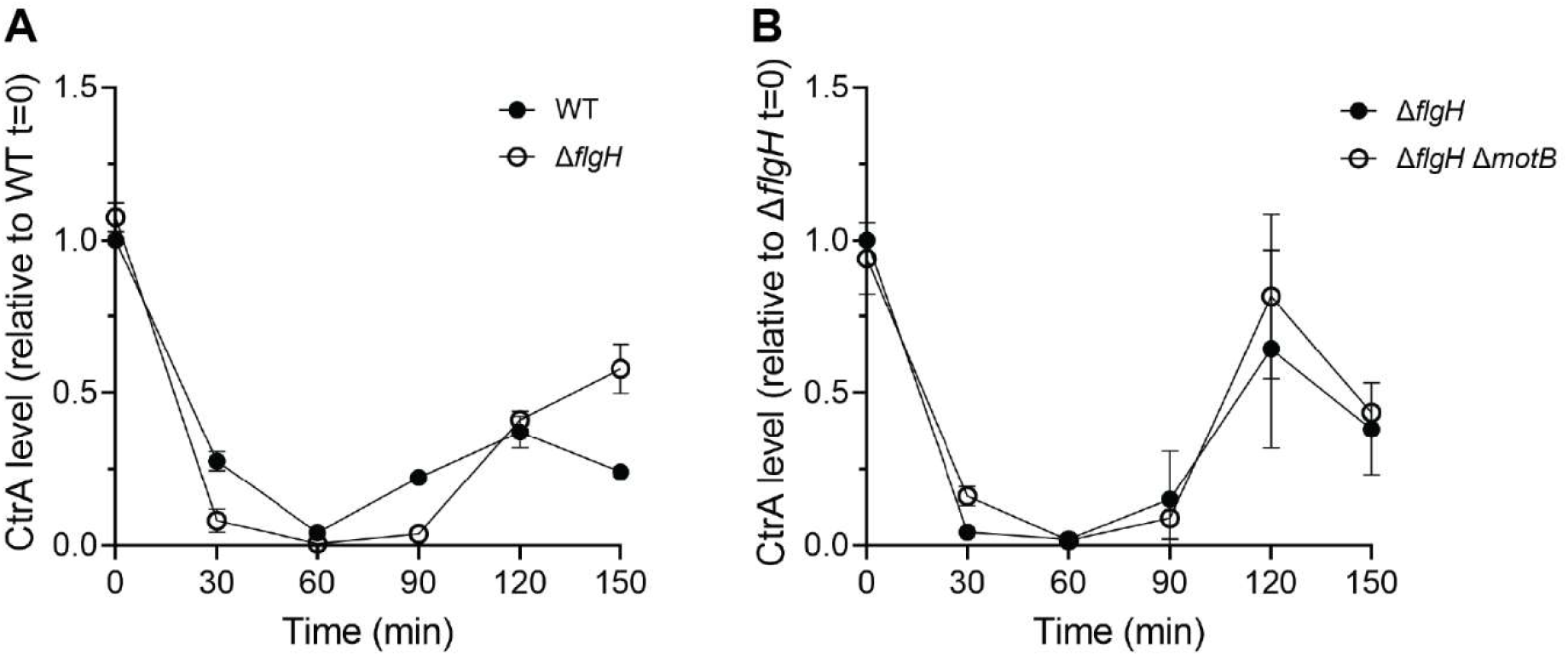
CtrA levels in synchronized WT, Δ*flgH*, and Δ*flgH* Δ*motB* cells. A) The Δ*flgH* strain has earlier and more sustained depletion of CtrA compared to WT. B) CtrA levels in the Δ*flgH* mutant do not change with the deletion of *motB*, indicating that CtrA depletion is not the mechanistic basis of the Δ*flgH* mutant’s earlier entry into S phase.

**Fig. S9:**
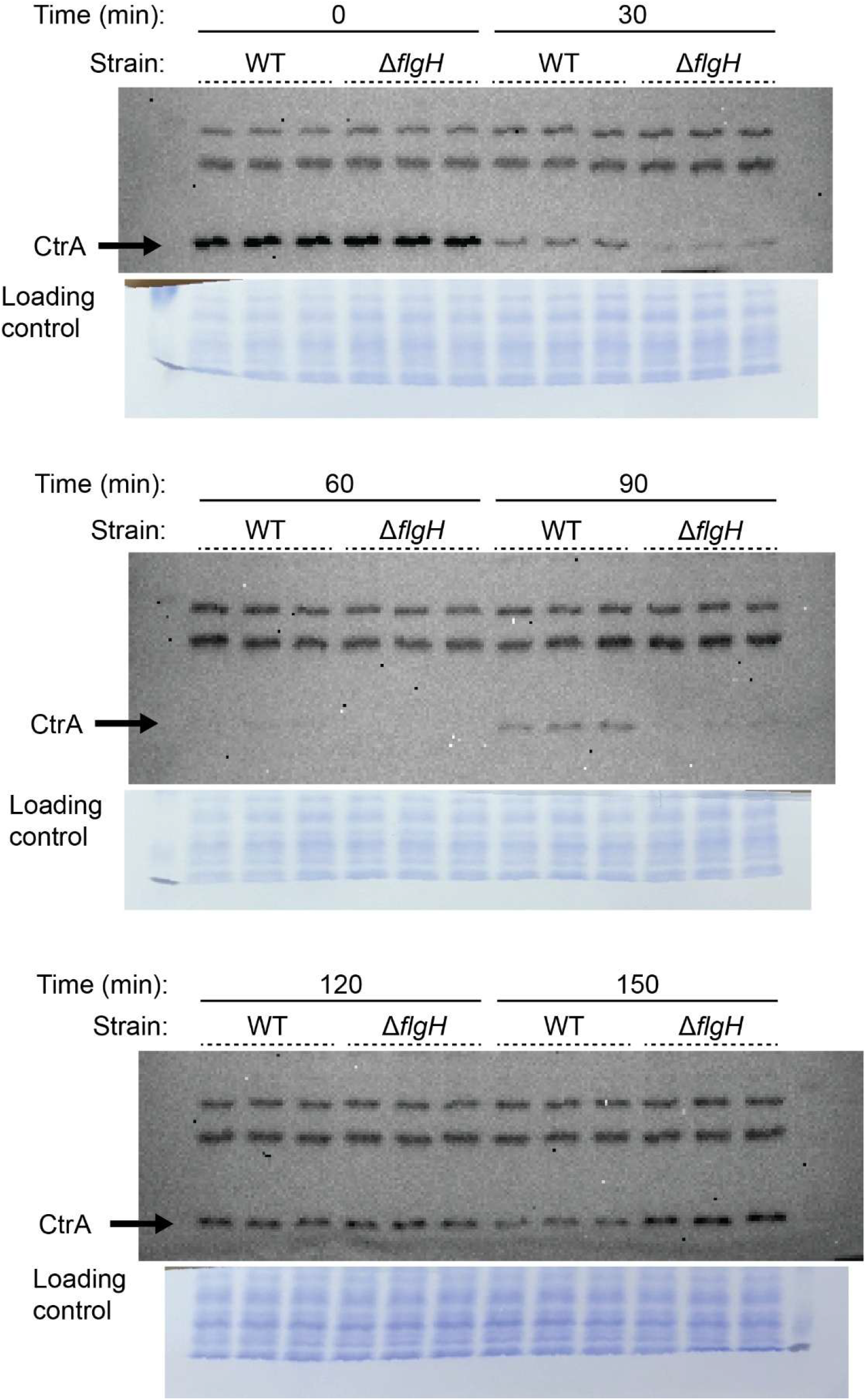
Immunoblot corresponding to Figure S8A.

**Fig. S10:**
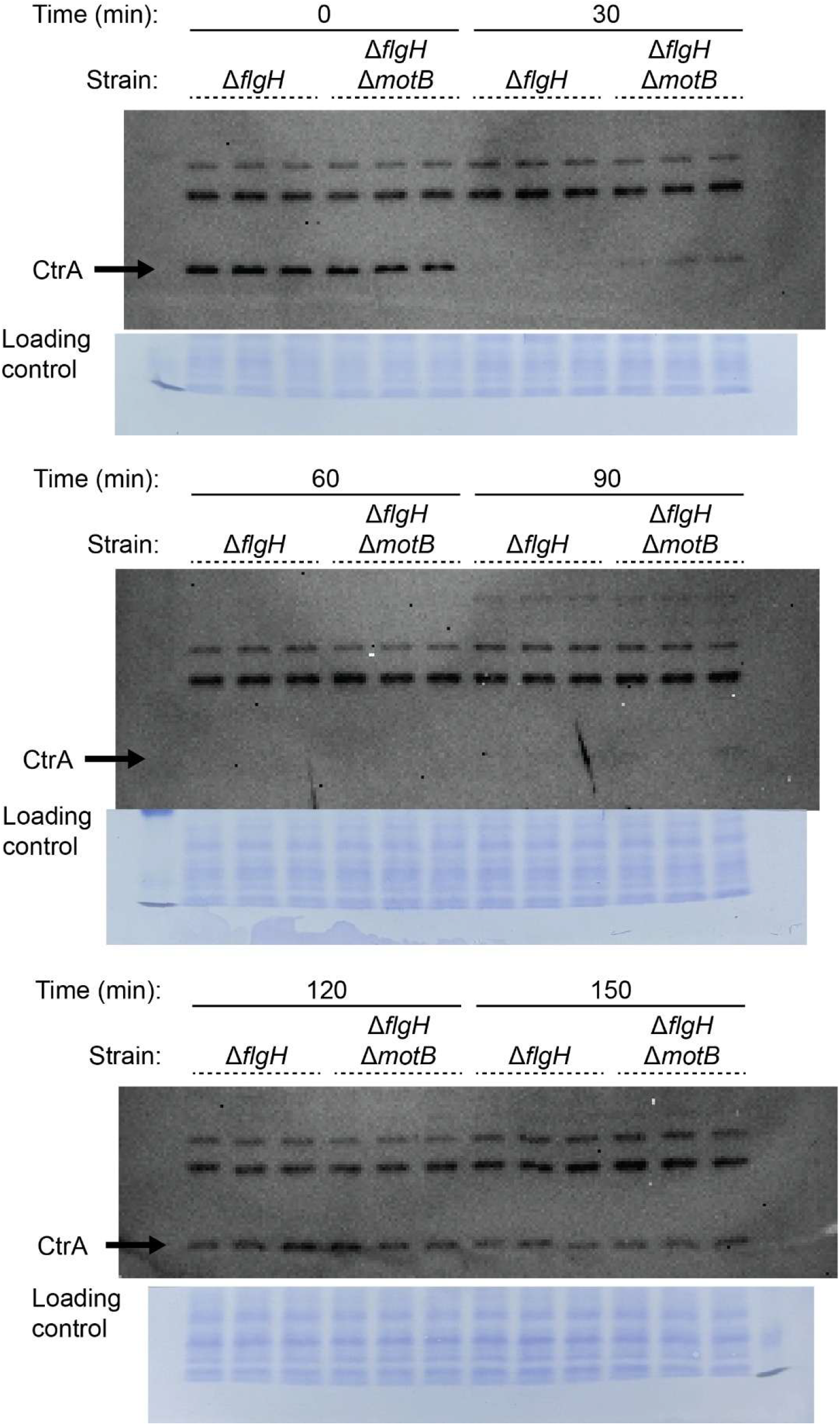
Immunoblot corresponding to Figure S8B

---

**Table S1:** Plasmids used in this study.

| Plasmid | Description | Antibiotic | Reference |
|---|---|---|---|
| pNPTS138 | Suicide plasmid for making unmarked deletions in *C. crescentus;* carries *sacB* for counter-selection | Km | M. R. Alley |
| pDH758 | pMT770, pXCTCY-2, integrating vector for xylose inducible expression of proteins in Caulobacter | Km | (69) |
| pDH1161 | To fuse fluorescent mVenus protein to C-terminus of PleC; Gibson cloning of *mvenus* fused between 3’ end and downstream region of CC_2482 | Km | (46) |
| pDH1209 | To fuse fluorescent mKate2 protein to C-terminus of DivJ; Gibson cloning of *mkate2* fused between 3’ end and downstream region of CC_ 1063 | Km | (46) |
| pDH1335 | To fuse fluorescent mVenus protein to N-terminus of MipZ; Gibson cloning of *mvenus* fused between upstream region and 5’ end of CC_2165 | Km | (46) |
| pDH1380 | To fuse fluorescent mVenus protein to C-terminus of FliG; Gibson cloning of *mvenus* fused between 3’ end and downstream region of CC_ 0906 | Km | (47) |
| pDH1615 | To fuse fluorescent mVenus protein to C-terminus of BacA; Gibson cloning of *mvenus* fused between 3’ end and downstream region of CC_1873 | Km | This study |
| pDH1616 | To fuse fluorescent mVenus protein to C-terminus of PleD; Gibson cloning of *mvenus* fused between 3’ end and downstream region of CC_2462 | Km | This study |
| pDH418 | To delete *flgH*; Gibson cloning of fused upstream and downstream regions of CC_2066 | Km | (45) |
| pDH805 | To delete *fliF;* Gibson cloning of fused upstream and downstream regions of CC_0905 | Km | (18) |
| pDH804 | To delete *fssC*; Gibson cloning of fused upstream and downstream regions of CC_2008 | Km | (46) |
| pDH766 | To delete *fssA*; Gibson cloning of fused upstream and downstream regions of CC_1064 | Km | (18) |
| pDH835 | To delete *fssB*; Gibson cloning of fused upstream and downstream regions of CC_2058 | Km | (18) |
| pDH767 | To delete *fssD*; Gibson cloning of fused upstream and downstream regions of CC_3471 | Km | This study |
| pDH795 | To delete *fssE*; Gibson cloning of fused upstream and downstream regions of CC_3472 | Km | This study |
| pDH1617 | To delete *fssDE* operon; Gibson cloning of fused upstream and downstream regions of CC_3471 and CC_3472 | Km | This study |
| pDH1114 | To delete *fssF*; Gibson cloning of fused upstream and downstream regions of CC_2175 | Km | (47) |
| pDH644 | To delete *pleD*; Gibson cloning of fused upstream and downstream regions of CC_2462 | Km | (18) |
| pDH765 | To delete *motB*; Gibson cloning of fused upstream and downstream regions of CC_1573 | Km | (18) |
| pDH769 | To delete *dgcB*; Gibson cloning of fused upstream and downstream regions of CC_1850 | Km | (18) |
| pDH1618 | To delete *tacA*; Gibson cloning of fused upstream and downstream regions of CC_3315 | Km | This study |
| pDH1619 | To integrate gene encoding cdGreen2.1 at the *xyl* locus; Fused to *syn2* promoter and *rrnB* terminator | Km | This study |
| pDH1620 | To integrate *P<sub>fssD</sub>-empty* at the *xyl* locus | Km | This study |
| pDH1621 | To integrate *P<sub>fssD</sub>-fssD* at the *xyl* locus | Km | This study |
| pDH1622 | To integrate *P<sub>fssD</sub>-fssDE* at the *xyl* locus | Km | This study |
| pDH1623 | To integrate *P<sub>fssD</sub>-fssE* at the *xyl* locus | Km | This study |

**Table S2:** Strains used in this study.

| **Strain** | **Organism** | **Genotype** | **Description** | **Source** |
|---|---|---|---|---|
| DH103 | *C. crescentus* CB15 | CB15 | Wild-type | ATCC 19089 |
| DH1077 | *C. crescentus* NA1000 | NA1000 | Wild-type | J. Poindexter |
| DH783 | *C. crescentus* CB15 | Δ*fssA* | In frame deletion of CC_1064 | (18) |
| DH845 | *C. crescentus* CB15 | Δ*fssB* | In frame deletion of CC_2058 | (18) |
| DH817 | *C. crescentus* CB15 | Δ*fssC* | In frame deletion of CC_2008 | (46) |
| DH785 | *C. crescentus* CB15 | Δ*fssD* | In frame deletion of CC_3471 | This study |
| DH813 | *C. crescentus* CB15 | Δ*fssE* | In frame deletion of CC_3472 | This study |
| DH1624 | *C. crescentus* CB15 | Δ*fssDE* | In frame deletion of CC_3471 and CC_3472 | This study |
| DH1124 | *C. crescentus* CB15 | Δ*fssF* | In frame deletion of CC_2175 | (47) |
| DH659 | *C. crescentus* CB15 | Δ*pleD* | In frame deletion of CC_2462 | (18) |
| DH776 | *C. crescentus* CB15 | Δ*motB* | In frame deletion of CC_1573 | (18) |
| DH787 | *C. crescentus* CB15 | Δ*dgcB* | In frame deletion of CC_1850 | (18) |
| DH1625 | *C. crescentus* CB15 | Δ*pleD* Δ*fssDE* | In frame deletion of CC_2462, CC_3471, and CC_3472 | This study |
| DH1626 | *C. crescentus* CB15 | Δ*tacA* | In frame deletion of CC_3315 | This study |
| DH820 | *C. crescentus* CB15 | Δ*fssC* Δ*motB* | In frame deletion of CC_1573 and CC_2008 | This study |
| DH816 | *C. crescentus* CB15 | Δ*fliF* | In frame deletion of CC_0905 | (18) |
| DH875 | *C. crescentus* CB15 | Δ*fliF* Δ*fssA* | In frame deletion of CC_0905 and CC_1064 | (18) |
| DH878 | *C. crescentus* CB15 | Δ*fliF* Δ*fssB* | In frame deletion of CC_0905 and CC_2058 | (18) |
| DH1154 | *C. crescentus* CB15 | Δ*fliF* Δ*fssC* | In frame deletion of CC_0905 and CC_2008 | (46) |
| DH1002 | *C. crescentus* CB15 | Δ*fliF* Δ*fssD* | In frame deletion of CC_0905 and CC_3471 | This study |
| DH1034 | *C. crescentus* CB15 | Δ*fliF* Δ*fssE* | In frame deletion of CC_0905 and CC_3472 | This study |
| DH1129 | *C. crescentus* CB15 | Δ*fliF* Δ*fssF* | In frame deletion of CC_0905 and CC_2175 | (47) |
| DH843 | *C. crescentus* CB15 | Δ*fliF* Δ*pleD* | In frame deletion of CC_0905 and CC_2462 | (18) |
| DH844 | *C. crescentus* CB15 | Δ*fliF* Δ*motB* | In frame deletion of CC_0905 and CC_1573 | (18) |
| DH876 | *C. crescentus* CB15 | Δ*fliF* Δ*dgcB* | In frame deletion of CC_0905 and CC_1850 | (18) |
| DH1627 | *C. crescentus* CB15 | Δ*fliF* Δ*fssDE* | In frame deletion of CC_0905, CC_3471, and CC3472 | This study |
| DH1628 | *C. crescentus* CB15 | Δ*fliF* Δ*pleD* Δ*fssDE* | In frame deletion of CC_0905, CC_2462, CC_3471, and CC3472 | This study |
| DH1629 | *C. crescentus* CB15 | Δ*fliF* Δ*tacA* | In frame deletion of CC_3315 and CC_0905 | This study |
| DH553 | *C. crescentus* CB15 | Δ*flgH* | In frame deletion of CC_2066 | (45) |
| DH784 | *C. crescentus* CB15 | Δ*flgH* Δ*fssA* | In frame deletion of CC_2066 and CC_1064 | (18) |
| DH846 | *C. crescentus* CB15 | Δ*flgH* Δ*fssB* | In frame deletion of CC_2066 and CC_2058 | (18) |
| DH818 | *C. crescentus* CB15 | Δ*flgH* Δ*fssC* | In frame deletion of CC_2066 and CC_2008 | (46) |
| DH786 | *C. crescentus* CB15 | Δ*flgH* Δ*fssD* | In frame deletion of CC_2066 and CC_3471 | This study |
| DH1043 | *C. crescentus* CB15 | Δ*flgH* Δ*fssE* | In frame deletion of CC_2066 and CC_3472 | This study |
| DH1125 | *C. crescentus* CB15 | Δ*flgH* Δ*fssF* | In frame deletion of CC_2066 and CC_2175 | (47) |
| DH649 | *C. crescentus* CB15 | Δ*flgH* Δ*pleD* | In frame deletion of CC_2066 and CC_2462 | (18) |
| DH777 | *C. crescentus* CB15 | Δ*flgH* Δ*motB* | In frame deletion of CC_2066 and CC_1573 | (18) |
| DH788 | *C. crescentus* CB15 | Δ*flgH* Δ*dgcB* | In frame deletion of CC_2066 and CC_1850 | (18) |
| DH1630 | *C. crescentus* CB15 | Δ*flgH* Δ*fssDE* | In frame deletion of CC_2066, CC_3471, and CC_3472 | This study |
| DH1631 | *C. crescentus* CB15 | Δ*flgH* Δ*pleD* Δ*fssDE* | In frame deletion of CC_2066, CC_2462, CC_3471, and CC_3472 | This study |
| DH1632 | *C. crescentus* CB15 | Δ*flgH* Δ*tacA* | In frame deletion of CC_3315 and CC_2066 | This study |
| DH1633 | *C. crescentus* CB15 | Δ*flgH* Δ*fssC* Δ*motB* | In frame deletion of CC_2066, CC_2008, and CC_1573 | This study |
| DH1634 | *C. crescentus* NA1000 | Δ*fssA* | In frame deletion of CCNA_01117 | This study |
| DH1635 | *C. crescentus* NA1000 | Δ*fssB* | In frame deletion of CCNA_02137 | This study |
| DH1338 | *C. crescentus* NA1000 | Δ*fssC* | In frame deletion of CCNA_02087 | (46) |
| DH1636 | *C. crescentus* NA1000 | Δ*fssD* | In frame deletion of CCNA_03585 | This study |
| DH1637 | *C. crescentus* NA1000 | Δ*fssE* | In frame deletion of CCNA_03586 | This study |
| DH1638 | *C. crescentus* NA1000 | Δ*fssF* | In frame deletion of CCNA_02257 | This study |
| DH1639 | *C. crescentus* NA1000 | Δ*pleD* | In frame deletion of CCNA_02546 | This study |
| DH1640 | *C. crescentus* NA1000 | Δ*motB* | In frame deletion of CCNA_01644 | This study |
| DH1641 | *C. crescentus* NA1000 | Δ*dgcB* | In frame deletion of CCNA_01926 | This study |
| DH1424 | *C. crescentus* NA1000 | Δ*flgH* | In frame deletion of CCNA_02145 | This study |
| DH1642 | *C. crescentus* NA1000 | Δ*flgH* Δ*motB* | In frame deletion of CCNA_02145 and CCNA_01644 | This study |
| DH1210 | *C. crescentus* CB15 | *pleC*::*pleC-venus* *divJ*::*divJ-mkate2* | Replacement of *divJ* with *divJ-mkate2* and *pleC* with *pleC-venus* in DH103 background | (46) |
| DH1643 | *C. crescentus* CB15 | Δ*fssA* *pleC*::*pleC-venus* *divJ*::*divJ-mkate2* | In frame deletion of CC_1064 in DH1210 background | This study |
| DH1644 | *C. crescentus* CB15 | Δ*fssB* *pleC*::*pleC-venus* *divJ*::*divJ-mkate2* | In frame deletion of CC_2058 in DH1210 background | This study |
| DH1219 | *C. crescentus* CB15 | Δ*fssC* *pleC*::*pleC-venus* *divJ*::*divJ-mkate2* | In frame deletion of CC_2008 in DH1210 background | (46) |
| DH1645 | *C. crescentus* CB15 | Δ*fssD* *pleC*::*pleC-venus* *divJ*::*divJ-mkate2* | In frame deletion of CC_3471 in DH1210 background | This study |
| DH1646 | *C. crescentus* CB15 | Δ*fssE* *pleC*::*pleC-venus* *divJ*::*divJ-mkate2* | In frame deletion of CC_3472 in DH1210 background | This study |
| DH1647 | *C. crescentus* CB15 | Δ*fssF* *pleC*::*pleC-venus* *divJ*::*divJ-mkate2* | In frame deletion of CC_2175 in DH1210 background | This study |
| DH1648 | *C. crescentus* CB15 | Δ*pleD* *pleC*::*pleC-venus* *divJ*::*divJ-mkate2* | In frame deletion of CC_2462 in DH1210 background | This study |
| DH1649 | *C. crescentus* CB15 | Δ*motB* *pleC*::*pleC-venus* *divJ*::*divJ-mkate2* | In frame deletion of CC_1573 in DH1210 background | This study |
| DH1650 | *C. crescentus* CB15 | Δ*dgcB* *pleC*::*pleC-venus* *divJ*::*divJ-mkate2* | In frame deletion of CC_1850 in DH1210 background | This study |
| DH1651 | *C. crescentus* CB15 | Δ*fssC* Δ*motB* *pleC*::*pleC-venus* *divJ*::*divJ-mkate2* | In frame deletion of CC_1573 in DH1219 background | This study |
| DH1425 | *C. crescentus* CB15 | *mipZ*::*venus-mipZ* | Replacement of *mipZ* with *venus-mipZ* in DH103 background | This study |
| DH1652 | *C. crescentus* CB15 | Δ*fssC* *mipZ*::*venus-mipZ* | Replacement of *mipZ* with *venus-mipZ* in DH817 background | This study |
| DH1653 | *C. crescentus* CB15 | Δ*fssD* *mipZ*::*venus-mipZ* | Replacement of *mipZ* with *venus-mipZ* in DH785 background | This study |
| DH1654 | *C. crescentus* CB15 | Δ*fssE* *mipZ*::*venus-mipZ* | Replacement of *mipZ* with *venus-mipZ* in DH813 background | This study |
| DH1655 | *C. crescentus* CB15 | Δ*fssF* *mipZ*::*venus-mipZ* | Replacement of *mipZ* with *venus-mipZ* in DH1124 background | This study |
| DH1656 | *C. crescentus* CB15 | Δ*pleD* *mipZ*::*venus-mipZ* | Replacement of *mipZ* with *venus-mipZ* in DH659 background | This study |
| DH1657 | *C. crescentus* CB15 | Δ*motB* *mipZ*::*venus-mipZ* | Replacement of *mipZ* with *venus-mipZ* in DH776 background | This study |
| DH1658 | *C. crescentus* CB15 | Δ*dgcB* *mipZ*::*venus-mipZ* | Replacement of *mipZ* with *venus-mipZ* in DH787 background | This study |
| DH1659 | *C. crescentus* CB15 | Δ*flgH* *mipZ*::*venus-mipZ* | Replacement of *mipZ* with *venus-mipZ* in DH513 background | This study |
| DH1660 | *C. crescentus* CB15 | Δ*flgH* Δ*fssC* *mipZ*::*venus-mipZ* | Replacement of *mipZ* with *venus-mipZ* in DH818 background | This study |
| DH1661 | *C. crescentus* CB15 | Δ*flgH* Δ*motB* *mipZ*::*venus-mipZ* | Replacement of *mipZ* with *venus-mipZ* in DH777 background | This study |
| DH1662 | *C. crescentus* CB15 | Δ*flgH* Δ*dgcB* *mipZ*::*venus-mipZ* | Replacement of *mipZ* with *venus-mipZ* in DH788 background | This study |
| DH1663 | *C. crescentus* CB15 | Δ*flgH* Δ*pleD* *mipZ*::*venus-mipZ* | Replacement of *mipZ* with *venus-mipZ* in DH649 background | This study |
| DH1664 | *C. crescentus* CB15 | Δ*fssC* Δ*motB* *mipZ*::*venus-mipZ* | Replacement of *mipZ* with *venus-mipZ* in DH820 background | This study |
| DH1665 | *C. crescentus* CB15 | Δ*flgH* Δ*fssC* Δ*motB* *mipZ*::*venus-mipZ* | Replacement of *mipZ* with *venus-mipZ* in DH1633 background | This study |
| DH1355 | *C. crescentus* CB15 | *fliG*::*fliG-venus* | Replacement of *fliG* with *fliG-venus* in DH103 background | (47) |
| DH1666 | *C. crescentus* CB15 | Δ*fssA* *fliG*::*fliG-venus* | Replacement of *fliG* with *fliG-venus* in DH783 background | This study |
| DH1667 | *C. crescentus* CB15 | Δ*fssB* *fliG*::*fliG-venus* | Replacement of *fliG* with *fliG-venus* in DH845 background | This study |
| DH1668 | *C. crescentus* CB15 | Δ*fssC* *fliG*::*fliG-venus* | Replacement of *fliG* with *fliG-venus* in DH817 background | This study |
| DH1669 | *C. crescentus* CB15 | Δ*fssD* *fliG*::*fliG-venus* | Replacement of *fliG* with *fliG-venus* in DH785 background | This study |
| DH1670 | *C. crescentus* CB15 | Δ*fssE* *fliG*::*fliG-venus* | Replacement of *fliG* with *fliG-venus* in DH813 background | This study |
| DH1364 | *C. crescentus* CB15 | Δ*fssF* *fliG*::*fliG-venus* | Replacement of *fliG* with *fliG-venus* in DH1124 background | (47) |
| DH1671 | *C. crescentus* CB15 | Δ*pleD* *fliG*::*fliG-venus* | Replacement of *fliG* with *fliG-venus* in DH659 background | This study |
| DH1672 | *C. crescentus* CB15 | Δ*motB* *fliG*::*fliG-venus* | Replacement of *fliG* with *fliG-venus* in DH776 background | This study |
| DH1673 | *C. crescentus* CB15 | Δ*dgcB* *fliG*::*fliG-venus* | Replacement of *fliG* with *fliG-venus* in DH787 background | This study |
| DH1674 | *C. crescentus* CB15 | Δ*flgH* *fliG*::*fliG-venus* | Replacement of *fliG* with *fliG-venus* in DH513 background | This study |
| DH1675 | *C. crescentus* CB15 | Δ*fssDE* *fliG*::*fliG-venus* | Replacement of *fliG* with *fliG-venus* in DH1624 background | This study |
| DH1676 | *C. crescentus* CB15 | Δ*pleD* Δ*fssDE* *fliG*::*fliG-venus* | Replacement of *fliG* with *fliG-venus* in DH1625 background | This study |
| DH1677 | *C. crescentus* CB15 | *bacA*::*bacA-venus* | Replacement of *bacA* with *bacA-venus* in DH103 background | This study |
| DH1678 | *C. crescentus* CB15 | Δ*fssC* *bacA*::*bacA-venus* | Replacement of *bacA* with *bacA-venus* in DH817 background | This study |
| DH1679 | *C. crescentus* CB15 | Δ*fssD* *bacA*::*bacA-venus* | Replacement of *bacA* with *bacA-venus* in DH785 background | This study |
| DH1680 | *C. crescentus* CB15 | Δ*fssE* *bacA*::*bacA-venus* | Replacement of *bacA* with *bacA-venus* in DH813 background | This study |
| DH1681 | *C. crescentus* CB15 | Δ*fssF* *bacA*::*bacA-venus* | Replacement of *bacA* with *bacA-venus* in DH1124 background | This study |
| DH1682 | *C. crescentus* CB15 | Δ*pleD* *bacA*::*bacA-venus* | Replacement of *bacA* with *bacA-venus* in DH659 background | This study |
| DH1683 | *C. crescentus* CB15 | Δ*motB* *bacA*::*bacA-venus* | Replacement of *bacA* with *bacA-venus* in DH776 background | This study |
| DH1684 | *C. crescentus* CB15 | Δ*dgcB* *bacA*::*bacA-venus* | Replacement of *bacA* with *bacA-venus* in DH787 background | This study |
| DH1685 | *C. crescentus* CB15 | *pleD*::*pleD-venus* | Replacement of *pleD* with *pleD-venus* in DH103 background | This study |
| DH1686 | *C. crescentus* CB15 | Δ*flgH* *pleD*::*pleD-venus* | Replacement of *pleD* with *pleD-venus* in DH513 background | This study |
| DH1687 | *C. crescentus* CB15 | Δ*fssDE* *pleD*::*pleD-venus* | Replacement of *pleD* with *pleD-venus* in DH1624 background | This study |
| DH1688 | *C. crescentus* CB15 | Δ*flgH* Δ*fssDE* *pleD*::*pleD-venus* | Replacement of *pleD* with *pleD-venus* in DH1630 background | This study |
| DH1689 | *C. crescentus* CB15 | *xyl*::*Psyn2-cdGreen2.1-rrnBterm* | Integration of pDH1619 into DH103 background | This study |
| DH1690 | *C. crescentus* CB15 | Δ*fssD* *xyl*::*Psyn2-cdGreen2.1-rrnBterm* | Integration of pDH1619 into DH785 background | This study |
| DH1691 | *C. crescentus* CB15 | Δ*fssE* *xyl*::*Psyn2-cdGreen2.1-rrnBterm* | Integration of pDH1619 into DH815 background | This study |
| DH1692 | *C. crescentus* CB15 | Δ*fssDE* *xyl*::*Psyn2-cdGreen2.1-rrnBterm* | Integration of pDH1619 into DH1624 background | This study |
| DH1693 | *C. crescentus* CB15 | Δ*pleD* *xyl*::*Psyn2-cdGreen2.1-rrnBterm* | Integration of pDH1619 into DH659 background | This study |
| DH1694 | *C. crescentus* CB15 | Δ*pleD* Δ*fssDE* *xyl*::*Psyn2-cdGreen2.1-rrnBterm* | Integration of pDH1619 into DH1625 background | This study |
| DH1695 | *C. crescentus* CB15 | Δ*flgH* *xyl*::*Psyn2-cdGreen2.1-rrnBterm* | Integration of pDH1619 into DH553 background | This study |
| DH1696 | *C. crescentus* CB15 | Δ*flgH* Δ*fssD* *xyl*::*Psyn2-cdGreen2.1-rrnBterm* | Integration of pDH1619 into DH786 background | This study |
| DH1697 | *C. crescentus* CB15 | Δ*flgH* Δ*fssE* *xyl*::*Psyn2-cdGreen2.1-rrnBterm* | Integration of pDH1619 into DH1043 background | This study |
| DH1698 | *C. crescentus* CB15 | Δ*flgH* Δ*fssDE* *xyl*::*Psyn2-cdGreen2.1-rrnBterm* | Integration of pDH1619 into DH1630 background | This study |
| DH1699 | *C. crescentus* CB15 | Δ*flgH* Δ*pleD* *xyl*::*Psyn2-cdGreen2.1-rrnBterm* | Integration of pDH1619 into DH649 background | This study |
| DH1700 | *C. crescentus* CB15 | Δ*flgH* Δ*pleD* Δ*fssDE* *xyl*::*Psyn2-cdGreen2.1-rrnBterm* | Integration of pDH1619 into DH1631 background | This study |
| DH1701 | *C. crescentus* CB15 | Δ*fssDE* *pleC*::*pleC-venus* *divJ*::*divJ-mkate2* | In frame deletion of CC_3471 and CC3472 in DH1210 background | This study |
| DH1702 | *C. crescentus* CB15 | Δ*pleD* Δ*fssDE* *pleC*::*pleC-venus* *divJ*::*divJ-mkate2* | In frame deletion of CC_2462 in DH1701 background | This study |
| DH1703 | *C. crescentus* CB15 | Δ*dgcB* Δ*fssDE* | In frame deletion of CC_3471, CC_3472, and CC_1850 | This study |
| DH1000 | *C. crescentus* CB15 | Δ*dgcB* Δ*pleD* | In frame deletion of CC_1850 and CC_2462 | (18) |
| DH1704 | *C. crescentus* CB15 | Δ*flgH* Δ*dgcB* Δ*fssDE* | In frame deletion of CC_2066, CC_3471, CC_3472, and CC_1850 | This study |
| DH1377 | *C. crescentus* CB15 | Δ*flgH* Δ*dgcB* Δ *pleD* | In frame deletion of CC_2066, CC_1850, and CC_2462 | (47) |
| DH1336 | *C. crescentus* NA1000 | *mipZ*::*venus-mipZ* | Replacement of *mipZ* with *venus-mipZ* in DH1077 background | (46) |
| DH1337 | *C. crescentus* NA1000 | Δ*fssC* *mipZ*::*venus-mipZ* | Replacement of *mipZ* with *venus-mipZ* in DH1338 background | (46) |
| DH1705 | *C. crescentus* NA1000 | Δ*motB* *mipZ*::*venus-mipZ* | Replacement of *mipZ* with *venus-mipZ* in DH1640 background | This study |
| DH1706 | *C. crescentus* CB15 | Δ*motB* Δ*pleD* *pleC*::*pleC-venus* *divJ*::*divJ-mkate2* | In frame deletion of CC_1573 and CC_2462 in DH1210 background | This study |
| DH1707 | *C. crescentus* CB15 | Δ*fssC* Δ*pleD* *pleC*::*pleC-venus* *divJ*::*divJ-mkate2* | In frame deletion of CC_2008 and CC_2462 in DH1210 background | This study |
| DH1708 | *C. crescentus* CB15 | Δ*flgH* Δ*fssD* *xyl*::*P<sub>fssD</sub>-empty* | Integration of pDH1620 into DH786 | This study |
| DH1709 | *C. crescentus* CB15 | Δ*flgH* Δ*fssD* *xyl*::*P<sub>fssD</sub>-fssD* | Integration of pDH1621 into DH786 | This study |
| DH1710 | *C. crescentus* CB15 | Δ*flgH* Δ*fssD* *xyl*::*P<sub>fssD</sub>-fssE* | Integration of pDH1623 into DH786 | This study |
| DH1711 | *C. crescentus* CB15 | Δ*flgH* Δ*fssD* *xyl*::*P<sub>fssD</sub>-fssDE* | Integration of pDH1622 into DH786 | This study |
| DH1712 | *C. crescentus* CB15 | Δ*flgH* Δ*fssE* *xyl*::*P<sub>fssD</sub>-empty* | Integration of pDH1620 into DH1043 | This study |
| DH1713 | *C. crescentus* CB15 | Δ*flgH* Δ*fssE* *xyl*::*P<sub>fssD</sub>-fssD* | Integration of pDH1621 into DH1043 | This study |
| DH1714 | *C. crescentus* CB15 | Δ*flgH* Δ*fssE* *xyl*::*P<sub>fssD</sub>-fssE* | Integration of pDH1623 into DH1043 | This study |
| DH1715 | *C. crescentus* CB15 | Δ*flgH* Δ*fssE* *xyl*::*P<sub>fssD</sub>-fssDE* | Integration of pDH1622 into DH1043 | This study |

## References

1. Vani S, Vadakkan K, Mani B. 2023. A narrative review on bacterial biofilm: its formation, clinical aspects and inhibition strategies. Futur J Pharm Sci 9:50.

2. Sharma S, Mohler J, Mahajan SD, Schwartz SA, Bruggemann L, Aalinkeel R. 2023. Microbial Biofilm: A Review on Formation, Infection, Antibiotic Resistance, Control Measures, and Innovative Treatment. Microorganisms 11:1614.

3. Li G, Brown PJB, Tang JX, Xu J, Quardokus EM, Fuqua C, Brun YV. 2012. Surface contact stimulates the just-in-time deployment of bacterial adhesins. Mol Microbiol 83:41–51.

4. McCarter L, Hilmen M, Silverman M. 1988. Flagellar dynamometer controls swarmer cell differentiation of V. parahaemolyticus. Cell 54:345–351.

5. Hug I, Deshpande S, Sprecher KS, Pfohl T, Jenal U. 2017. Second messenger–mediated tactile response by a bacterial rotary motor. Science 358:531–534.

6. Belas R. 2014. Biofilms, flagella, and mechanosensing of surfaces by bacteria. Trends in Microbiology 22:517–527.

7. Laventie B-J, Jenal U. 2020. Surface Sensing and Adaptation in Bacteria. Annual Review of Microbiology 74:735–760.

8. Del Medico L, Cerletti D, Schächle P, Christen M, Christen B. 2020. The type IV pilin PilA couples surface attachment and cell-cycle initiation in Caulobacter crescentus. Proceedings of the National Academy of Sciences 117:9546–9553.

9. Snyder RA, Ellison CK, Severin GB, Whitfield GB, Waters CM, Brun YV. 2020. Surface sensing stimulates cellular differentiation in Caulobacter crescentus. Proceedings of the National Academy of Sciences 117:17984–17991.

10. Berne C, Ellison CK, Ducret A, Brun YV. 2018. Bacterial adhesion at the single-cell level. Nat Rev Microbiol 16:616–627.

11. Kuchma SL, Geiger CJ, Webster SS, Fu Y, Montoya R, O’Toole GA. 2025. Genetic analysis of flagellar-mediated surface sensing by Pseudomonas aeruginosa PA14. Journal of Bacteriology 207:e00520–24.

12. Cairns LS, Marlow VL, Bissett E, Ostrowski A, Stanley-Wall NR. 2013. A mechanical signal transmitted by the flagellum controls signalling in Bacillus subtilis. Molecular Microbiology 90:6–21.

13. Obeid Charrouf F, Whitfield GB, Ellison CK, Brun YV. 2025. Stimulation of the Caulobacter crescentus surface sensing pathway by deletion of a specialized minor pilin-like gene. mBio 16:e02302–25.

14. Ardissone S, Viollier PH. 2015. Interplay between flagellation and cell cycle control in *Caulobacter*. Current Opinion in Microbiology 28:83–92.

15. Minamino T, Kinoshita M. Structure, Assembly, and Function of Flagella Responsible for Bacterial Locomotion. EcoSal Plus 11:eesp-0011-2023.

16. Stock D, Namba K, Lee LK. 2012. Nanorotors and self-assembling macromolecular machines: The torque ring of the bacterial flagellar motor. Current Opinion in Biotechnology 23:545–554.

17. Santiveri M, Roa-Eguiara A, Kühne C, Wadhwa N, Hu H, Berg HC, Erhardt M, Taylor NMI. 2020. Structure and Function of Stator Units of the Bacterial Flagellar Motor. Cell 183:244–257.e16.

18. Hershey DM, Fiebig A, Crosson S. 2021. Flagellar Perturbations Activate Adhesion through Two Distinct Pathways in Caulobacter crescentus. mBio 12:e03266–20.

19. Ellison CK, Kan J, Dillard RS, Kysela DT, Ducret A, Berne C, Hampton CM, Ke Z, Wright ER, Biais N, Dalia AB, Brun YV. 2017. Obstruction of pilus retraction stimulates bacterial surface sensing. Science 358:535–538.

20. Römling U, Galperin MY, Gomelsky M. 2013. Cyclic di-GMP: the First 25 Years of a Universal Bacterial Second Messenger. Microbiol Mol Biol Rev 77:1–52.

21. Boyd CD, O’Toole GA. 2012. Second Messenger Regulation of Biofilm Formation: Breakthroughs in Understanding c-di-GMP Effector Systems. Annu Rev Cell Dev Biol 28:439–462.

22. Valentini M, Filloux A. 2016. Biofilms and Cyclic di-GMP (c-di-GMP) Signaling: Lessons from *Pseudomonas aeruginosa* and Other Bacteria*. Journal of Biological Chemistry 291:12547–12555.

23. O’Malley MR, Dearing HN, Zheng X, Kretschmer AN, Cho TH-S, Raivio TL, Parsek MR. 2025. The Pseudomonas aeruginosa Cpx system provides a cyclic-di-GMP independent link between cell envelope stress and surface sensing. mBio 17:e02726–25.

24. Holland LM, O’Donnell ST, Ryjenkov DA, Gomelsky L, Slater SR, Fey PD, Gomelsky M, O’Gara JP. 2008. A staphylococcal GGDEF domain protein regulates biofilm formation independently of cyclic dimeric GMP. J Bacteriol 190:5178–5189.

25. Govers SK, Jacobs-Wagner C. 2020. Caulobacter crescentus: model system extraordinaire. Curr Biol 30:R1151–R1158.

26. Shapiro L, Agabian-Keshishian N, Bendis I. 1971. Bacterial Differentiation. Science 173:884–892.

27. Poindexter JS. 1964. BIOLOGICAL PROPERTIES AND CLASSIFICATION OF THE CAULOBACTER GROUP. Bacteriol Rev 28:231–295.

28. Curtis PD, Brun YV. 2010. Getting in the Loop: Regulation of Development in Caulobacter crescentus. Microbiology and Molecular Biology Reviews 74:13–41.

29. Wheeler RT, Shapiro L. 1999. Differential Localization of Two Histidine Kinases Controlling Bacterial Cell Differentiation. Molecular Cell 4:683–694.

30. Hecht GB, Newton A. 1995. Identification of a novel response regulator required for the swarmer-to-stalked-cell transition in Caulobacter crescentus. J Bacteriol 177:6223–6229.

31. Paul R, Weiser S, Amiot NC, Chan C, Schirmer T, Giese B, Jenal U. 2004. Cell cycle-dependent dynamic localization of a bacterial response regulator with a novel di-guanylate cyclase output domain. Genes Dev 18:715–727.

32. Kaczmarczyk A, Hempel AM, von Arx C, Böhm R, Dubey BN, Nesper J, Schirmer T, Hiller S, Jenal U. 2020. Precise timing of transcription by c-di-GMP coordinates cell cycle and morphogenesis in Caulobacter. Nat Commun 11:816.

33. Biondi EG, Skerker JM, Arif M, Prasol MS, Perchuk BS, Laub MT. 2006. A phosphorelay system controls stalk biogenesis during cell cycle progression in Caulobacter crescentus. Mol Microbiol 59:386–401.

34. Aldridge P, Paul R, Goymer P, Rainey P, Jenal U. 2003. Role of the GGDEF regulator PleD in polar development of Caulobacter crescentus. Molecular Microbiology 47:1695–1708.

35. Aldridge P, Jenal U. 1999. Cell cycle-dependent degradation of a flagellar motor component requires a novel-type response regulator. Molecular Microbiology 32:379–391.

36. Sommer JM, Newton A. 1989. Turning off flagellum rotation requires the pleiotropic gene pleD: pleA, pleC, and pleD define two morphogenic pathways in Caulobacter crescentus. Journal of Bacteriology 171:392–401.

37. Lori C, Ozaki S, Steiner S, Böhm R, Abel S, Dubey BN, Schirmer T, Hiller S, Jenal U. 2015. Cyclic di-GMP acts as a cell cycle oscillator to drive chromosome replication. Nature 523:236–239.

38. Laub MT, Chen SL, Shapiro L, McAdams HH. 2002. Genes directly controlled by CtrA, a master regulator of the Caulobacter cell cycle. Proc Natl Acad Sci U S A 99:4632–4637.

39. Domian IJ, Quon KC, Shapiro L. 1997. Cell Type-Specific Phosphorylation and Proteolysis of a Transcriptional Regulator Controls the G1-to-S Transition in a Bacterial Cell Cycle. Cell 90:415–424.

40. Quon KC, Yang B, Domian IJ, Shapiro L, Marczynski GT. 1998. Negative control of bacterial DNA replication by a cell cycle regulatory protein that binds at the chromosome origin. Proc Natl Acad Sci U S A 95:120–125.

41. Bastedo DP, Marczynski GT. 2009. CtrA response regulator binding to the Caulobacter chromosome replication origin is required during nutrient and antibiotic stress as well as during cell cycle progression. Molecular Microbiology 72:139–154.

42. Bodenmiller D, Toh E, Brun YV. 2004. Development of Surface Adhesion in Caulobacter crescentus. Journal of Bacteriology 186:1438–1447.

43. Tsang PH, Li G, Brun YV, Freund LB, Tang JX. 2006. Adhesion of single bacterial cells in the micronewton range. Proceedings of the National Academy of Sciences 103:5764–5768.

44. Hershey DM, Porfírio S, Black I, Jaehrig B, Heiss C, Azadi P, Fiebig A, Crosson S. 2019. Composition of the Holdfast Polysaccharide from Caulobacter crescentus. J Bacteriol 201:e00276–19.

45. Hershey DM, Fiebig A, Crosson S. 2019. A Genome-Wide Analysis of Adhesion in Caulobacter crescentus Identifies New Regulatory and Biosynthetic Components for Holdfast Assembly. mBio 10:e02273–18.

46. Hellenbrand CN, Stevenson DM, Gromek KA, Amador-Noguez D, Hershey DM. 2025. A deoxynucleoside triphosphate triphosphohydrolase promotes cell cycle progression in Caulobacter crescentus. Journal of Bacteriology 207:e00145–25.

47. Salemi RI, Cruz AK, Hershey DM. 2024. A flagellar accessory protein links chemotaxis to surface sensing. Journal of Bacteriology 206:e00404–24.

48. Gonzalez D, Collier J. 2014. Effects of (p)ppGpp on the Progression of the Cell Cycle of Caulobacter crescentus. Journal of Bacteriology 196:2514–2525.

49. Burton GJ, Hecht GB, Newton A. 1997. Roles of the histidine protein kinase pleC in Caulobacter crescentus motility and chemotaxis. J Bacteriol 179:5849–5853.

50. Kühn J, Briegel A, Mörschel E, Kahnt J, Leser K, Wick S, Jensen GJ, Thanbichler M. 2010. Bactofilins, a ubiquitous class of cytoskeletal proteins mediating polar localization of a cell wall synthase in Caulobacter crescentus. EMBO J 29:327–339.

51. Barrows JM, Goley ED. Synchronized Swarmers and Sticky Stalks: Caulobacter crescentus as a Model for Bacterial Cell Biology. J Bacteriol 205:e00384–22.

52. Kaczmarczyk A, van Vliet S, Jakob RP, Teixeira RD, Scheidat I, Reinders A, Klotz A, Maier T, Jenal U. 2024. A genetically encoded biosensor to monitor dynamic changes of c-di-GMP with high temporal resolution. Nat Commun 15:3920.

53. Thanbichler M, Shapiro L. 2006. MipZ, a spatial regulator coordinating chromosome segregation with cell division in Caulobacter. Cell 126:147–162.

54. Sobol G, Hershey DM. 2026. The gac system integrates physical and chemical cues to promote plant root attachment. Applied and Environmental Microbiology 92:e00778–26.

55. Pijper A. 1947. Methylcellulose and Bacterial Motility. Journal of Bacteriology 53:257–269.

56. Flemming H-C, Wuertz S. 2019. Bacteria and archaea on Earth and their abundance in biofilms. Nat Rev Microbiol 17:247–260.

57. Carrascosa C, Raheem D, Ramos F, Saraiva A, Raposo A. 2021. Microbial Biofilms in the Food Industry—A Comprehensive Review. Int J Environ Res Public Health 18:2014.

58. Li Y, Qu Y, Yang H, Zhou X, Xiao P, Shao T. 2023. Combatting biofilms in potable water systems: A comprehensive overview to ensuring industrial water safety. Environmental Microbiology Reports 15:445–454.

59. Mishra S, Huang Y, Li J, Wu X, Zhou Z, Lei Q, Bhatt P, Chen S. 2022. Biofilm-mediated bioremediation is a powerful tool for the removal of environmental pollutants. Chemosphere 294:133609.

60. Perry EK, Tan M-W. 2023. Bacterial biofilms in the human body: prevalence and impacts on health and disease. Front Cell Infect Microbiol 13:1237164.

61. Chen JC, Viollier PH, Shapiro L. 2005. A membrane metalloprotease participates in the sequential degradation of a Caulobacter polarity determinant. Molecular Microbiology 55:1085–1103.

62. Curtis PD, Quardokus EM, Lawler ML, Guo X, Klein D, Chen JC, Arnold RJ, Brun YV. 2012. The scaffolding and signaling functions of a localization factor impact polar development. Mol Microbiol 84:712–735.

63. Abel S, Bucher T, Nicollier M, Hug I, Kaever V, Abel zur Wiesch P, Jenal U. 2013. Bi-modal Distribution of the Second Messenger c-di-GMP Controls Cell Fate and Asymmetry during the Caulobacter Cell Cycle. PLoS Genet 9:e1003744.

64. Abel S, Chien P, Wassmann P, Schirmer T, Kaever V, Laub MT, Baker TA, Jenal U. 2011. Regulatory Cohesion of Cell Cycle and Cell Differentiation through Interlinked Phosphorylation and Second Messenger Networks. Molecular Cell 43:550–560.

65. Laventie B-J, Sangermani M, Estermann F, Manfredi P, Planes R, Hug I, Jaeger T, Meunier E, Broz P, Jenal U. 2019. A Surface-Induced Asymmetric Program Promotes Tissue Colonization by *Pseudomonas aeruginosa*. Cell Host & Microbe 25:140–152.e6.

66. Schneider CA, Rasband WS, Eliceiri KW. 2012. NIH Image to ImageJ: 25 years of image analysis. Nat Methods 9:671–675.

67. Ducret A, Quardokus EM, Brun YV. 2016. MicrobeJ, a tool for high throughput bacterial cell detection and quantitative analysis. Nat Microbiol 1:16077.

68. Quon KC, Marczynski GT, Shapiro L. 1996. Cell cycle control by an essential bacterial two-component signal transduction protein. Cell 84:83–93.

69. Thanbichler M, Iniesta AA, Shapiro L. 2007. A comprehensive set of plasmids for vanillate- and xylose-inducible gene expression in Caulobacter crescentus. Nucleic Acids Res 35:e137.

